# Ecology and molecular targets of hypermutation in the global microbiome

**DOI:** 10.1101/2020.04.01.020958

**Authors:** Simon Roux, Blair G. Paul, Sarah C. Bagby, Michelle A. Allen, Graeme Attwood, Ricardo Cavicchioli, Ludmila Chistoserdova, Steven J. Hallam, Maria E. Hernandez, Matthias Hess, Wen-Tso Liu, Michelle A. O’Malley, Xuefeng Peng, Virginia I. Rich, Scott Saleska, Emiley A. Eloe-Fadrosh

**Affiliations:** DOE Joint Genome Institute, Berkeley, CA; Marine Biological Laboratory, Woods Hole, MA; Case Western Reserve University, Cleveland, OH; The University of New South Wales, Sydney, New South Wales, Australia; AgResearch Limited, Grasslands Research Centre, Palmerston North, New Zealand; University of Washington, Seattle, WA; Department of Microbiology & Immunology, University of British Columbia, Vancouver, Canada; Graduate Program in Bioinformatics, University of British Columbia, Genome Sciences Centre, Vancouver, Canada; Genome Science and Technology Program, University of British Columbia, Vancouver, Canada; Life Sciences Institute, University of British Columbia, Vancouver, Canada; ECOSCOPE Training Program, University of British Columbia, Vancouver, Canada; Instituto de Ecología A.C. Red de Manejo Biotechnológico de Recursos. Xalapa, Veracruz, México; University of California Davis, Davis, CA; University of Illinois at Urbana-Champaign, Urbana, IL; Department of Chemical Engineering, University of California Santa Barbara, Santa Barbara, CA; Marine Science Institute, University of California Santa Barbara, Santa Barbara, CA; Ohio State University, Columbus, OH; University of Arizona, Tucson, AZ

**Author notes:** Correspondence to &.

## Abstract

Changes in the sequence of an organism’s genome, i.e. mutations, are the raw material of evolution^1^. The frequency and location of mutations can be constrained by specific molecular mechanisms, such as Diversity-generating retroelements (DGRs)^2–4^. DGRs introduce mutations in specific target genes, and were characterized from several cultivated bacteria and bacteriophages^2^. Whilst a larger diversity of DGR loci has been identified in genomic data from environmental samples, i.e. metagenomes, the ecological role of these DGRs and their associated evolutionary drivers remain poorly understood^5–7^. Here we built and analyzed an extensive dataset of >30,000 metagenome-derived DGRs, and determine that DGRs have a single evolutionary origin and a universal bias towards adenine mutations. We further identified six major lineages of DGRs, each associated with a specific ecological niche defined as a genome type, i.e. whether the DGR is encoded on a viral or cellular genome, a limited set of taxa and environments, and a distinct type of target. Finally, we leverage read mapping and metagenomic time series to demonstrate that DGRs are consistently and broadly active, and responsible for >10% of all amino acid changes in some organisms at a conservative estimate. Overall, these results highlight the strong constraints under which DGRs diversify and expand, and elucidate several distinct roles these elements play in natural communities and in shaping microbial community structure and function in our environment.

## Introduction

Diversity-generating retroelements (DGRs) are genetic elements that can produce a large number of mutations in a specific region of a target gene^2,4^. The first DGR identified induces hypervariation in a structural protein responsible for host recognition and attachment of bacteriophage BPP-1^3^. Other examples of DGRs were subsequently characterized, with the best-studied instances, in *Legionella* and *Treponema*, targeting surface-displayed proteins^8,9^. All currently known DGRs seem to use the same molecular mechanism, known as mutagenic retrohoming, to generate hypervariation in the target protein^4,10,11^. Mechanistically, a DGR requires three main components: a reverse transcriptase (RT); a template region (TR), which in most cases is intergenic; and a variable region (VR), that is near-identical to the TR and located within the coding sequence of the target protein. The DGR-encoded RT uses a primary TR transcript as the template for error-prone reverse transcription. Our current understanding is that DGR RTs are strongly promiscuous at template adenines, leading to the incorporation of dATP, dGTP, and dCTP in roughly equal proportions to the templated dTTP. The resulting sequence variant, i.e. TR-cDNA, is then integrated in the protein-coding gene, replacing the original VR sequence, through a yet-undefined homing mechanism^10,11^.

Building on the handful of well-characterized DGRs, recent studies have sought to explore DGR diversity by mining genomic data for DGR-like RT genes found next to imperfect repeats with mismatches opposing adenine positions. This approach was successfully applied to both draft genomes^2,12^ and metagenome assemblies^5–7,13–17^. Collectively, these studies identified ∼1,500 DGRs, and suggested that DGRs are present in diverse environments ranging from the human gut to deep-sea sediments and terrestrial groundwater^5,6,13^. DGRs were also associated with a broad range of genomes including uncultivated bacteria from the Candidate Phyla Radiation (CPR) and several archaea^5,6^.

While DGRs are now broadly recognized as important diversification mechanisms in microbes, their specific activity and role(s) across organisms and biomes remain elusive. Specifically, ecological and evolutionary drivers of targeted hypermutation are currently unknown due to the lack of a global contextualized map of DGRs. Similarly, predicting the potential role of individual DGRs is currently challenging because the vast majority of putative targets are functionally uncharacterized. Therefore, it remains unclear for which proteins, functions, organisms, or environments this type of targeted hyperdiversification constitutes a selective advantage.

Here we expand the set of known DGRs ∼15-fold by extracting 31,007 DGRs from public metagenomes and metatranscriptomes to obtain a holistic view of DGR diversity and their spatial and temporal dynamics. We leverage this comprehensive collection to (i) evaluate the global ecology and evolution of DGRs across viral and cellular genomes, (ii) characterize the functional diversity and molecular constraints of DGR targets, and (iii) infer temporal patterns of DGR activity across organisms and biomes. Taken together, these analyses reveal how DGRs are frequently transferred between genomes yet clearly restricted to specific ecological niches, within which they likely impact both viral and microbial dynamics by consistently driving amino acid-level diversification of their target domains.

## Results

### Large-scale metagenome mining uncovers an extensive diversity of DGRs

To identify candidate DGRs, we searched for reverse transcriptase (RT) genes found within 1kb of an imperfect repeat, and used phylogenetic placement and mismatch patterns to identify false-positive detections (see Methods). We applied this approach to 81,404 public genomes and 9,467 public metagenomes to obtain a global view of DGR diversity (Supplementary Table 1). In genomes, we detected a total of 1,314 DGRs, comparable in number and diversity to those identified in previous mining of genome databases^2,6^. By contrast, we detected 31,007 DGRs in public metagenomes, a ∼15-fold increase compared to the total number of DGRs previously reported^5–7,13–17^. Overall, DGRs were detected from ≥1,500 bacterial and archaeal genera and ≥90 environment types (Supplementary Table 2, Supplementary Text).

In order to systematically explore this DGR diversity, we used average amino acid identity (AAI) to group RT sequences, first into 13,415 OTUs (≥95% AAI), then into 1,318 clusters (≥50% AAI, Supplementary Fig. S1, Supplementary Tables 3 & 4). Members of each OTU and cluster were associated with consistent genome (i.e., viral vs cellular), taxa, and biome types, suggesting that these groupings represent distinct DGR evolutionary units (Supplementary Text, Supplementary Fig. S2). To evaluate global DGR diversity, a phylogenetic tree including a representative of each RT clusters was built (Fig. 1A). DGRs formed a monophyletic clade separated from other types of RTs, such as introns or retrons, supporting a single evolutionary origin for these elements^2^. Overall, 75% of clusters were composed exclusively of metagenome-derived DGR sequences, and our survey alone represented an almost 6-fold increase in DGR phylogenetic diversity (573%), highlighting the significant contribution of metagenome and metatranscriptome assemblies to the exploration of DGR sequence space.

**Figure 1.**
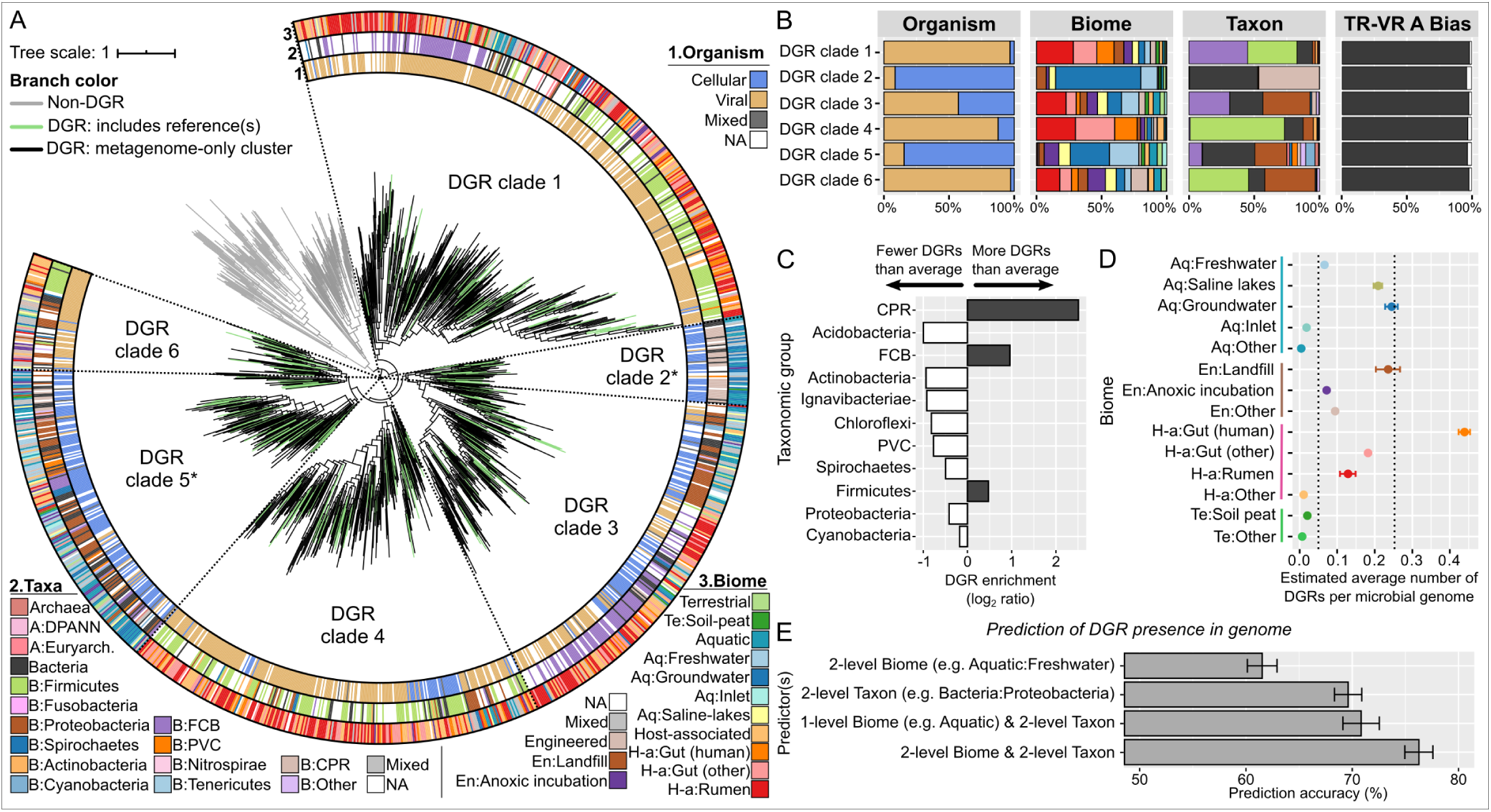
Distribution of DGR diversity across organisms, biomes, and taxa. A. Phylogeny of DGR and non-DGR reverse transcriptases (RT). RT protein sequences were first grouped into “RT clusters”, and a representative for each cluster was included in the tree building process (see Methods). Branches are colored according to the type of RT in the corresponding cluster. All nodes with support <50% were collapsed. From inside to outside, the outer rings display the consensus genome type, taxonomic classification, and biome of each RT cluster. CPR: Candidate Phyla Radiation. DPANN: *Diapherotrites, Parvarchaeota, Aenigmarchaeota, Nanoarchaeota, Nanohaloarchaeota*. FCB: *Flavobacteria, Fibrobacteres, Chlorobi, Bacteroides*. PVC: *Planctomycetes, Verrucomicrobia, Chlamydiae*. Aq: Aquatic. Te: Terrestrial. En: Engineered. H-a: Host-associated. NA correspond to cases for which the feature could not be estimated (see Supplementary Fig. S2). B. Distribution of each feature at the RT OTU level across DGR clades. Colors used in the bar chart are identical to panel A, and NA values were not included. C. Enrichment of DGR-encoding genomes across taxa. The total number of genomes observed across metagenome assemblies was calculated based on single-copy marker genes (see Methods), and an average frequency of DGR was derived from the entire dataset. A frequency of DGR detection per genome was then calculated for each taxonomic group and compared to the overall frequency to derive log2 enrichment ratios. All log2-ratios presented in the figure are statistically significant (Chi-square test of independence corrected p-value <1E-10) except for the Cyanobacteria group (p-value=0.21). D. DGR enrichment across biomes. For each biome, a linear regression was computed between the estimated total number of genomes and the number of DGRs detected in each metagenome (see Supplementary Fig. S5). The regression slope was then considered as an estimation of the average number of DGR per genome and is displayed here with error bars representing the standard error of the slope estimation. Cutoffs of 0.05 and 0.2 DGRs per genome are highlighted with vertical dashed lines. For these calculations, viral and low complexity metagenomes were excluded. E. Accuracy of random forest classifiers trained to predict whether an input genome encodes a DGR. The features used as predictors are indicated on the y-axis, and all classifiers were trained and evaluated on the same dataset (see Methods). An interactive version of the tree is available at http://itol.embl.de/tree/15713110928194161585007032.

### DGRs dispersion is strongly constrained and reflected in cohesive lineage partitioning

Mapping the genome type, taxonomy, and biome of each DGR cluster onto the tree suggested that the global DGR diversity could be divided into 6 main clades (DGR clades 1-6, Fig. 1A & B, Supplementary Text). Three clades (DGR clades 1, 4, and 6) are composed of DGRs identified almost exclusively in viruses, predominantly phages infecting abundant gut bacteria belonging to the *Firmicutes* and *Bacteroidetes* (DGR clade 1), or *Proteobacteria* (DGR clades 4 & 6, Fig. 1B, Supplementary Fig. S3, Supplementary Text). Clades 2 and 5 are almost entirely composed of cellular-encoded DGRs, mostly from aquatic biomes, and either restricted to the *Patescibacteria*, referred to as the Candidate Phyla Radiation (CPR, DGR clade 2), or affiliated to diverse phyla including *Proteobacteria* and *Bacteroidetes* (DGR clade 5). Finally, clade 3 includes a nearly even mix of virus- and cell-derived DGRs, mostly associated with *Proteobacteria* and *Bacteroidetes*. Across all clades, alignments between template repeat (TR) and variable repeat (VR) overwhelmingly displayed ≥75% of mismatches facing adenine residues in the TR: after manual inspection of outliers (Supplementary Text), we identified only 7 clusters of seemingly genuine DGRs with <75% of mismatches facing adenine residues (Supplementary Fig. S4, Supplementary Table 4). This suggests that the mutation bias towards adenine is an intrinsic feature of DGR RTs. The monophyly in the RT tree and near-universality of the adenine mutation bias also suggests a single origin for all modern DGRs followed by sporadic transfers across organisms and biomes leading to the 6 main clades observed here (Supplementary Text).

Across the 9,467 metagenomes we examined, several taxa and biomes were clearly enriched in DGRs. DGRs were significantly more common (p-value <10^−16^) in members of the CPR, *Firmicutes*, and *Flavobacteria*-*Bacteroidetes*-*Chlorobi* (FCB) groups (Fig. 1C). Similarly, we observed a significantly higher rate of DGR detection per genome in several environments, including human gut microbiomes, saline lakes, landfills, and groundwater reservoirs (Fig. 1D, Supplementary Fig. S5). Notably, a random forest classifier was able to predict with >75% accuracy whether a genome encodes a DGR based simply on a 2-level biome classification (e.g. Host-Associated:Human gut) and a 2-level taxonomic classification (e.g. Bacteria:Cyanobacteria). The prediction accuracy decreased when only one of these two features was used, indicating that DGRs are associated with specific taxon-biome combinations (Fig. 1E). Taken together, these results point towards a long and complex DGR evolutionary history, with DGRs able to transfer between unrelated organisms, but only being selected for and retained in specific niches as evidenced by their current uneven distribution across genomes and biomes.

### DGR targets are diverse but share a conserved organization

To gain insight into the potential roles of identified DGRs, we next investigated the diversity of 36,611 putative DGR target genes present in genomes and metagenomes. As previously reported^2^, the majority (68%) of these targets could not be functionally annotated when individually compared to reference databases. However, *de novo* clustering revealed that most DGR targets (>92%) grouped into 24 protein clusters (PCs) representing just four broad functional classes (see below), and clearly partitioned by genome type and DGR clade (Fig. 2A, Supplementary Table 5, Supplementary Text).

**Figure 2.**
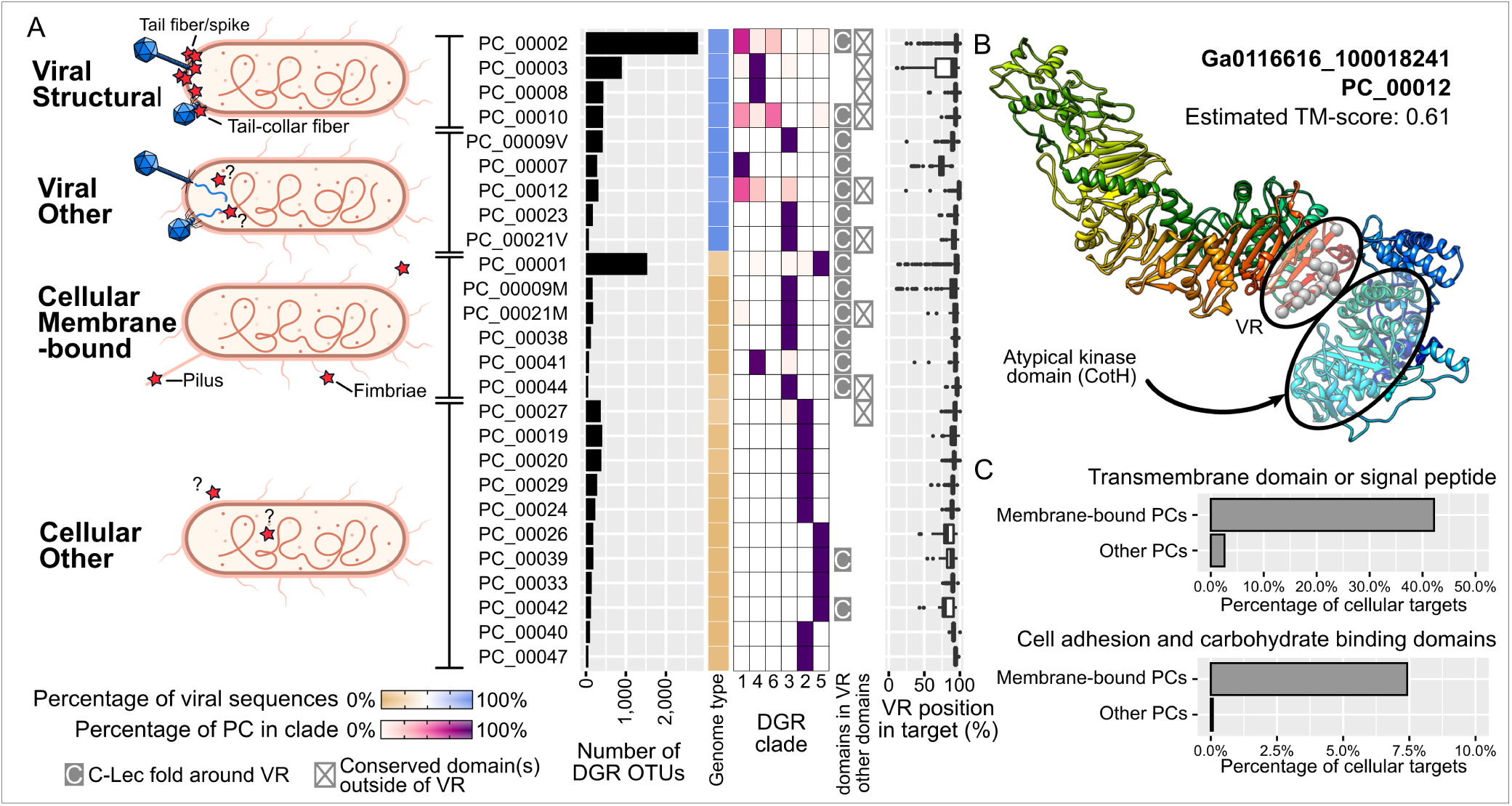
Diversity and major types of DGR targets. A. Prevalence and sequence characteristics of the most abundant DGR target protein clusters (PCs). The 24 PCs listed here represent >92% of all identified DGR targets. These were divided into four major types (left panel; Target proteins highlighted with red stars). Characteristics of each PC are indicated to the right, including number of associated DGR RT OTUs, relative proportion of viral-encoded vs cellular-encoded DGRs (“Genome type”), distribution across DGR clades, detection of a C-Lec fold around the VR region, detection of a conserved domain outside of the VR region, and relative position of the VR region within the target sequence. The boxplot lower and upper hinges correspond to the first and third quartiles, respectively, and the whiskers extend no further than ±1.5 times the interquartile range. B. Predicted structure of a viral-encoded target sequence from PC_00012 displaying similarity to a eukaryotic-like kinase domain (CotH). The structure is colored with a blue-red rainbow gradient from the N- to C-terminal end and predicted variable residues in the VR (i.e. corresponding to TR adenines) are highlighted with grey spheres. Because of the large size of the protein (2,284 aa), structure prediction was run on a subset of the sequence from position 786 to 2,284, i.e. without the N-terminal region. The model quality was assessed based on the TM-score estimated by I-TASSER (a TM-score >0.5 indicates a model with a likely correct topology). C. Percentage of cellular targets with a predicted transmembrane domain (top) or one or more functional domain(s) associated with cell adhesion and/or carbohydrate binding identified outside of the VR region (bottom). Target sequences are divided based on their PC membership into “membrane-bound” PCs or “other” PCs (see panel A).

Analysis of functional domains and residue conservation across PCs suggested a near-universal modular organization. Targets were typically multi-domain proteins, with the VR region found at the C-termini, as previously reported^2^ (Fig. 2A). While these C-terminal regions include DGR-variable residues, they were overall more conserved than the rest of the sequence across all PCs, likely due to structural constraints associated with DGR-induced hypervariation^18,19^ (Supplementary Fig. S6). Accordingly, whereas a range of folds were predicted for N-terminal domains, annotated VR-containing regions were systematically associated with C-type lectin (C-Lec) folds. Some rare VRs had previously been tentatively linked to Ig-like folds^2^, our analysis suggests these targets instead correspond to phage tail fibers containing Ig-like domain(s) next to an uncharacterized, non-Ig-like, VR domain (Supplementary Fig. S7, Supplementary Text). Since novel variants of C-Lec fold domains are still being discovered on a regular basis^18,20^, it is probable that other uncharacterized conserved domains overlapping VR regions represent new variants of the C-Lec fold domain family. Based on the distribution of corresponding target PCs, these novel C-Lec fold will most likely be associated with novel viruses and uncultivated bacteria (CPR) and archaea (Supplementary Fig. S8). The observed modularity of target proteins also suggests that intragenic recombination may occur for DGR targets, with the potential to fuse a wide range of independently folding domains to a C-terminal C-Lec-encoding region to produce a chimeric target ready for mutagenesis.

### DGR targets are primarily involved in virus-cell and cell-particle interactions

Given the near-universal modular organization of target proteins, putative functions were assigned based on the presence of conserved domains or sequence features identified outside of the C-terminal VR region. Viral targets from DGR Clades 1, 4, and 6 were found to mostly represent structural proteins, i.e. would be displayed on the surface of the virion and likely involved in host interaction (Fig. 2A). Since our knowledge of viral structural proteins is still partial^21^, target PCs currently lacking a functional annotation but associated with the same DGR clades (e.g. PC_00007) likely represent additional unknown structural proteins. Combined with the strong enrichment for viral-encoded DGRs in the mammalian gut, this suggests that mechanisms of host resistance to phages in these environments are skewed towards cell wall modification to circumvent host recognition and phage adsorption, which would select for hypervariation of phage structural proteins^22^. One notable exception was PC_00012, which included targets with an atypical eukaryotic-like serine/threonine kinase (CotH-like domain). While these kinase domains are typically located distantly from the VR, structural predictions suggested both regions may be in close contact, perhaps directly interacting with each other, once the protein is folded (Fig. 2B). Based on the known functions of eukaryotic-like serine/threonine kinases, these DGRs may be associated with the manipulation of fundamental host cell signaling pathways such as cell division, dormancy, or sporulation^23^.

For cellular targets, most PCs contained at least one N-terminal transmembrane domain or signal peptide, along with different functional domains involved in carbohydrate binding and cell adhesion (Fig. 2A & C, Supplementary Table 5). This suggests that most of these targets are membrane-anchored proteins binding extracellular substrates, potentially including particle aggregates and other microbial cells. Accordingly, metagenome-assembled genomes associated with the most prevalent of these targets (PC_00001) displayed a gene content and functional annotation consistent with a copiotrophic or particle-associated lifestyle (Supplementary Text). DGRs with PC_00001 targets were primarily detected in aquatic environments (Supplementary Fig. S8), however the frequency of DGR detection was highly variable between different aquatic biomes (Fig. 1, Supplementary Fig. S5). Taken together, this suggests that the selective advantage provided by broad-scale particle binding, cell-cell attachment, or surface adherence may vary between environments. For instance, in the open ocean, random binding may not be advantageous as it could lead to elevated cell loss due to sinking particles^24^, which may explain why DGRs are rarely detected in these samples (Fig. 1). Unlike most other cellular-encoded targets, the ones encoded by CPRs in clade 2 and archaea in clade 5 do not typically include any transmembrane or recognizable domain (Fig. 2C), as was previously reported^5,6^. Whether this is due to functional domains not being readily identified in these divergent genomes or to these proteins representing genuine non-membrane-bound DGR targets remains to be investigated. Overall, the large collection of targets identified in this study provides additional and strong evidence that DGRs are primarily linked to cell-particle, cell-cell, and virus-cell interactions, and in some rare cases may be involved in microbial cell regulation^6^.

### DGRs are broadly active across organisms and biomes

Next, we evaluated population diversity for DGR loci across taxa and ecosystems. To that end, we analyzed single nucleotide and amino acid variants for 6,901 DGRs with ≥10 kb genomic context and ≥20x coverage (see Methods). Overall, single-nucleotide variants (SNVs) could be detected for 70.1% of the VR loci (Fig. 3A, Supplementary Fig. S9, Supplementary Text). When SNVs were detected, VR loci were strongly enriched in non-synonymous SNVs, as estimated through pN/pS ratio (Fig. 3B): while non-target genes displayed a low frequency of non-synonymous SNVs and pN/pS ratios consistent with a purifying selection (i.e., <1), >80% of VR loci with ≥1 SNV(s) displayed pN/pS ratios >1, indicating a strong enrichment in non-synonymous mutations.

**Figure 3.**
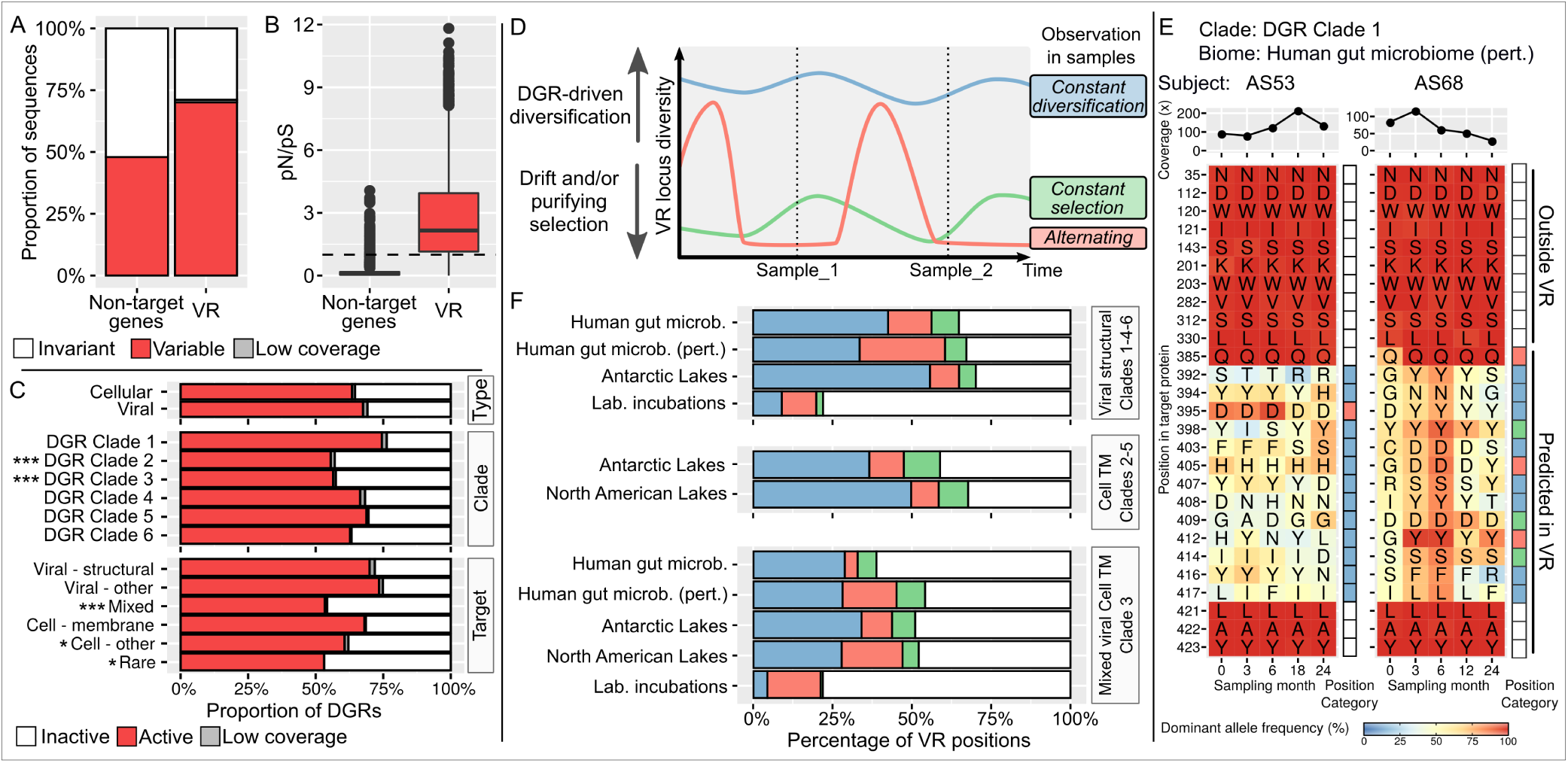
Diversity patterns associated with DGR target loci. A. Proportion of genes with ≥1 SNV observed (synonymous or non-synonymous), for both non-target genes within 10kb of a DGR RT (left), and VR loci (right). “Low coverage” category includes cases in which the coverage of the VR region was significantly lower than that of the surrounding genes, suggesting that the read recruitment may be only partial, and the population diversity in the VR can not be reliably inferred (see Supplementary Fig. S9). B. Distribution of pN/pS values for genes with ≥1 synonymous SNV, for both non-target genes and VR loci. A dashed line indicates pN/pS=1. Boxplot lower and upper hinges correspond to the first and third quartiles, whiskers extend no further than ±1.5*Inter-quartile range. C. Proportion of DGRs estimated “active” vs “inactive” based on an enrichment of VR loci in non-synonymous SNVs compared to surrounding genes, across different DGR classifications. Groups with a significantly lower proportion of active sequences (Chi-squared test of independence) are highlighted with star symbols (Bonferroni-corrected p-values: *<1E-03, **<1E-05, ***<1E-10). D. Schematic representation of the two competing forces exerted on VR loci: purifying selection and DGR diversification. Three examples of possible DGR activity levels are indicated in color, with the resulting observations across a time series (“Sample_1” and “Sample_2”) summarized in the right column. E. Example of diversity and changes observed for one DGR target across two time series datasets. For each position, the corresponding amino acid is indicated in the main heatmap with its frequency within the population indicated in color. The right panel indicates the category of the position based on within-sample entropy, between-samples cosine distances, and number of amino acid changes in the time series (see Supplementary Text), colored as in panel D. The top panel indicates the median coverage of all positions in each sample. For reference purposes, 10 random positions from the same protein outside of the predicted VR are included. F. Distribution of VR positions into “activity” categories (see panel D) across different biomes and clades. Cases with <50% of variable positions and <5% of amino acid changes were considered as “low DGR activity” and colored in white. Only groups for which ≥10 DGRs were available are included.

Using the enrichment in non-synonymous SNVs as marker for recent DGR activity, we next compared different types of DGRs. For all DGR groups, 50-75% of DGRs showed signs of recent activity (Fig. 3C). Viral-associated DGRs were linked to the highest activity level, while members of DGR clades 2 and 3 displayed a significantly lower-than-average activity level (Supplementary Fig. S10). However, strong purifying selection that would reduce population diversity may mask DGR activity in these single-sample variant analyses, with active DGRs having the potential to generate new variants that would almost instantaneously be purged from the population and thus evade detection.

### DGR activity drives frequent changes in target residues

Given widespread DGR activity, most VR loci can be expected to evolve under two antagonistic forces: DGR diversification and purifying selection. The relationship between these forces can be examined from time-series data so long as the adaptive value of different variants fluctuates over time (Fig. 3D, Supplementary Fig. S11). Specifically, we hypothesized DGR diversification to lead to high population diversity within each sample and changes in dominant allele between time points, while purifying selection would reduce population diversity within each sample. For alternating phases of diversification and purifying selection, we expect to observe a low population diversity within each sample but changes in the dominant allele between samples (Supplementary Fig. S12).

To test this hypothesis and shed light on the balance between DGR diversification and purifying selection in nature, we analyzed the subset of DGRs found across metagenomic time-series. We identified 130 longitudinal data sets containing 563 DGRs amenable to analysis, i.e. having ≥10kb genomic context, with coverage ≥10x, and detected at ≥2 time-points, Supplementary Table 6). Overall, a majority of predicted VR positions in these DGRs showed high diversity associated with frequent amino acid replacement, suggesting a high DGR activity overpowering purifying selection (Fig. 3E and Fig. 3F). This pattern was consistent across natural biomes but absent for *in vitro* microbiomes, i.e. laboratory incubations. There, both the observed diversity and amino acid change frequency were much lower for all types of DGRs, probably due to the population bottlenecks^25,26^, lack of various environmental stress, and/or shorter time frames of these experiments (Fig. 3F). Overall, viral-encoded DGRs targeting structural proteins systematically displayed a higher rate of amino acid turnover than DGRs of the cellular-encoded clades or “mixed” clade 3 from the same environment (Fig. 3F, Supplementary Fig. S13). Extrapolating from the average mutation rates observed here, we conservatively estimated that DGR-driven mutations would be responsible for 6-16% of all amino acid changes in an average viral genome, even though DGRs target only ∼0.1% of amino acid residues (see Supplementary text). Meanwhile, DGRs were identified in human gut samples from individuals following a 12-month weight-loss program^27^, and in samples taken from infants during gut microbiome establishment^28^. DGR activity in these ‘perturbed’ human gut microbiomes was thus compared to activity in ‘unperturbed’ human gut microbiomes, obtained from adult HMP subjects with multiple visits^29^. Overall, DGR activity seemed to be higher in perturbed vs non-perturbed human gut microbiomes, with a more pronounced increase in activity for cellular-encoded clade 3 DGRs (Fig. 3F, Supplementary Fig. S14). Taken together, this suggests that DGRs drive more steady changes in viral structural proteins through time compared to non-structural or cellular targets, and some of the latter may be more associated with adaptation during stress episodes. Whether this is due to a stronger control of DGR RT activity or a stronger selection exerted outside of stress episodes on targets remains to be determined.

## Conclusion

The extensive comparative analysis of metagenome-derived DGRs presented here reveals fundamental aspects of DGR biology as a key component of microbial and viral genome evolution. The near-universal conservation of the adenine mutation bias and the C-Lec fold in target proteins suggests that DGR RTs are mechanistically constrained in the type and location of mutations they can generate and, reciprocally, that C-Lec folds have a seemingly unique ability to accommodate massive sequence variation^18^. The strong DGR enrichment observed in select biomes and taxa likely reflects specific ecological conditions and lifestyles for which hypermutation is advantageous. For instance, in human gut microbiomes, the combination of high resource availability and frequent infections by a broad diversity of phages is expected to favor resistance through cell wall modification^30^, which would in turn select for DGR-encoding viruses. Finally, the widespread and seemingly constant DGR activity suggests that these elements are mostly used to maintain a high population diversity at target loci. DGRs may thus represent a key mechanism by which viruses and cellular microorganisms modulate adaptation and response patterns in an environment of constant and unpredictable change.

## Supplementary Figures

Supplementary Figure S1: Size distribution of the 100 largest RT OTUs (top panel) and RT Clusters (bottom panel). The bars are colored according to the presence/absence of reference sequences in the corresponding OTU/Cluster.

Supplementary Figure S2: Characteristics of DGR RT OTUs and Clusters. Each bar chart indicates the consistency of one feature (taxonomic classification, biome, or genome type) across members of a DGR RT OTU (A) or DGR RT Cluster (B). For OTUs with ≥2 members (i.e. non-singletons) with inconsistent values for a feature, a majority rule was applied: if a majority of an OTU members had the same value, this value was used for the OTU (“Mixed values: solved”). In case of tie (i.e. equal number of members associated to different features), the OTU feature was considered as unknown (“Mixed values: irreconcilable”). For Clusters, a similar approach was used with a 2/3rd majority rule. All Clusters for which ≥ 2/3rd of the members had the same value were considered as “Consistent value” and the value was assigned to the cluster. Cases in which the majority value in the cluster was associated with < 2/3rd of the members were considered as “Mixed values: irreconcilable”.

Supplementary Figure S3: Phylogeny of isolate and metagenome-binned genomes encoding one or more DGRs and associated with human gut samples. Nodes with support <50 were collapsed, and nodes with support ≥80 are noted with a black circle. For each genome, the different clades of DGRs detected in the genome is indicated next to the tree as a colored heatmap. The genome relative abundance is then indicated next to the heatmap: isolate genomes are highlighted with grey squares, genome bins ranked as one of the 5 most abundant genomes within a metagenome are highlighted with black squares.

Supplementary Figure S4: Examples of predicted DGRs with atypical (non-A) mutation bias. For each DGR, the clade, genome type, taxonomic classification, biome, and primary target affiliation are indicated when available. The genome maps are colored based on each predicted CDS functional annotation: the DGR reverse-transcriptase in red, target gene in green, other genes in blue, and “hypothetical protein” in grey.

Supplementary Figure S5: Link between estimated total number of genomes (x-axis) and number of DGRs detected (y-axis) for metagenomes across different biomes. For each biome, a linear regression line is indicated in color, with the 95% confidence interval outlined in gray. Zoomed plots are displayed on the right panel, and the zoomed-in region is highlighted with a dashed black square on the full plot on the left panel.

Supplementary Figure S6: Average residue conservation in predicted targets. A. Example of average residue conservation in 35-residues windows along the multiple alignment of PC_00008. An “extended” VR region (200 residues upstream and 20 residues downstream of the average predicted VR region) is highlighted in grey, which corresponds to the variable residues and the surrounding conserved domain. B. Distribution of residue conservation in “extended VR” and non-VR regions for the 24 largest target clusters. All distribution were significantly different (Kolmogorov–Smirnov test p-value <2E-16). The magnitude of the difference between VR and non-VR region is indicated through Cohen’s d effect size (star symbols on the x-axis). All target PCs showed a higher average conservation in VR compared to non-VR regions except for PC_00012, which is highlighted with a black circle. The boxplot lower and upper hinges correspond to the first and third quartiles, respectively, and the whiskers extend no further than ±1.5 times the interquartile range.

Supplementary Figure S7: Phylogeny and domain organization of target sequences clustered in PC_00003. Nodes with support <50 were collapsed, and nodes with support ≥80 are indicated with a black circle. For each sequence or clade, a schematic of the domain organization is indicated to the right of the tree, with a black line proportional to the sequence length, VR domains indicated with a red circle, and other domains indicated with colored rectangles. Monophyletic clades with a consistent domain organization were collapsed. The two clades of sequences displaying an internal VR region typically followed by one or several Ig-like folds in C-terminal are highlighted with a black dashed square. Reference sequences identified in the “Ig1” and “Ig2” domains are noted with a star symbol next to the sequence name.

Supplementary Figure S8: Taxonomic classification and biome of DGR OTUs associated with the 24 largest target PCs. The PCs are ordered according to the 4 main categories of targets, as on Fig. 2. Target PCs for which the VR region was not identified as a putative C-Lec fold are highlighted in red. The proportion of DGR OTUs associated with specific taxa (left) or biomes (right) was calculated independently for each target PC. White cells in the heatmap correspond to an absence of DGR for the corresponding taxa/biome and target PC combination. CPR: Candidate Phyla Radiation. FCB: Flavobacteria, Fibrobacteres, Chlorobi, Bacteroides.

Supplementary Figure S9: Read mapping and SNV calling on VR regions. A. Comparison of coverage between VR regions and non-target genes for individual TR-VR pairs. Only cases with coverage ≥20x are displayed, and both x- and y-axis are displayed as log10 scale. The 1-to-1 line is indicated in black. A lower bound for a 95% confidence interval was calculated from the average coverage of non-target genes from the same contig minus 2 standard deviations. If the VR coverage was below this cutoff, it was considered as significantly lower than expected, the TR-VR was colored in blue in this plot, and flagged as “low coverage” if no SNVs were detected in Fig. 3A. B. Comparison of SNV density for individual VR regions obtained from Mpileup (x-axis) vs Freebayes (y-axis). A linear regression curve is plotted in blue, and the associated equation is indicated on the plot (p-value <2e-16).

Supplementary Figure S10: Distribution of active-vs-inactive DGRs across genome type, clade, and targets, for different ranges of coverage. Groups (i.e. DGRs of the same genome type, DGR clade, or target) with a significantly lower proportion of active sequences compared to the average of the corresponding coverage category (Chi-squared test of independence) are highlighted with star symbols (Bonferroni-corrected p-values: *<1E-03, **<1E-05, ***<1E-10).

Supplementary Figure S11: Schematic of the different categories of positions defined based on population diversity across time series. Each example represents an individual position observed across 5 samples. The population diversity in each sample is represented as a heatmap, and the two metrics used to define the DGR activity categories are plotted underneath, either for each sample for the entropy, or between pairs of consecutive samples for the cosine similarity.

Supplementary Figure S12: Examples of DGR target positions with changes in dominant amino acid between samples and low diversity within sample (“Alternating” pattern in Fig. 3D). For each position (y-axis), the corresponding amino acid is indicated in the main heatmap with its frequency within the population indicated in color for each sample (x-axis). The right panel indicates the category of the position based on within-sample entropy, between-samples cosine distances, and number of amino acid changes in the time series (see Supplementary Text), colored as in Fig. 3D. The top panel indicates the median coverage of all positions in each sample. For reference purposes, 10 random positions from the same protein but outside of the predicted VR are included in the heatmap.

Supplementary Figure S13: Percentage of positions with ≥1 change(s) in dominant allele among positions considered as “Constant diversity” or “Alternating” for different types of DGR across major biomes. For each biome, the percentage in “Viral structural – Clades 1/4/6” DGRs was compared to the percentage in other DGR categories combined using a Chi-square test of independence. Groups with a significantly higher proportion of positions with ≥1 change(s) are highlighted with star symbols (Bonferroni-corrected p-values: *<1E-03, **<1E-05, ***<1E-10). † Counts for DGRs associated with viral structural proteins in temperate lakes are based on only 5 DGRs, while all other environments had > 10 DGRs associated with viral structural proteins.

Supplementary Figure S14: Comparison of DGR activity between perturbed and non-perturbed human microbiome samples. Left panels display the distribution of activity categories for VR positions between perturbed and non-perturbed human gut microbiome DGRs. The conditions under which each dataset was collected are indicated at the bottom of the figure. The right panel bar graph indicates the number of observations (i.e. total number of DGRs covered in at least 2 time points across all subjects) for each dataset. Panel A includes all relevant DGRs, while panel B and C include only viral- or cellular-encoded DGRs, respectively. Statistically significant comparisons (Chi-square of independence) are highlighted with star symbols (Bonferroni-corrected p-values: *<1E-03, **<1E-05, ***<1E-10).

## Supplementary Table

Supplementary Table 1: List of metagenomes mined for DGRs. Metagenomes are associated with samples, ecological category, sample type, and publication based on information from IMG^31^ and Gold^32^. The number of genomes in each assembly was estimated based on single-copy marker genes, while the ratio of bp in viral sequences among contigs of 10kb or more was derived from VirSorter detection of viral contigs (see Methods). For laboratory incubations, the biome listed corresponds to the biome of the original sample, when available.

Supplementary Table 2: List of DGRs detected in public genomes and metagenomes. For cases where the mutation bias of a DGR was atypical (<75% A) while the average bias of the corresponding OTU was typical (≥75% A), the DGR bias was replaced by the OTU one with the mention “(from OTU)”. For OTUs with “unknown” genome type, a predicted one based on the taxonomy, biome, and target of the corresponding OTU is indicated if available, noted with “(predicted)”. NA: information unknown and not available.

Supplementary Table 3: List of DGR OTUs with annotation. When inconsistent, OTU-level annotation was derived from a consensus annotation (majority rule) of OTU members (see Methods). This list does not include the 655 RT OTUs that corresponded to non-DGR references included in the tree. NA: information unknown and not available.

Supplementary Table 4: List of DGR clusters with annotation. Cluster-level annotation was derived from a consensus annotation (2/3rd majority) of cluster members (see Methods). A list of the 7 clusters of seemingly genuine DGRs with atypical mutation biases is provided in a separate tab. This list does not include the 175 RT clusters that corresponded to non-DGR references included in the tree. NA: information unknown and not available.

Supplementary Table 5: List of predicted DGR targets with annotation. For VR region position, the best hit between the TR and the target sequence was considered. NA: information unknown and not available.

Supplementary Table 6: List of longitudinal datasets used in the time-series analyses. These datasets were previously analyzed in refs. ^27–29,33–35^

## Methods

### Collection and annotation of Reverse-transcriptase (RT) and DGR reference sets

Reference sequences of RTs (DGRs and non-DGR) were collected from ref.^36^. Additional DGR references were obtained from ref.^2^. For the latter, corresponding TR/VR (template repeat and variable repeat) and target sequences were extracted from the supplementary html and fasta files (respectively) provided. Taxonomic classification of reference sequences was derived from NCBI database based on the genome identifier provided for each DGR. For DGRs not taxonomically classified as viruses, the genome sequence was downloaded from NCBI and VirSorter was used to identify whether the DGR RTs was encoded in an integrated provirus or metagenomic viral contig.

### Detection of DGRs in (meta)genomes

The overall detection pipeline consisted of 3 main steps: (i) identification of RT based on matches to HMM profiles using hmmsearch v3.2.1^37^, (ii) detection of repeats around the candidate RT using blastn v2.9.0+^38^ with option -word_size 8 -dust no -gapopen 6 -gapextend 2, and (iii) selection of putative DGRs, TRs, and target genes based on repeat patterns and RT length. Importantly, unlike existing tools^39^, this detection was agnostic to the variation between the repeats, because it did not require mismatches between repeats to be associated with adenine residues. The input sequences for this detection pipeline were (i) all genes predicted from IMG public genomes, and (ii) genes predicted on contigs ≥1kb and encoding ≥2 genes from IMG public metagenomes (Supplementary Table 1). The former was used to complete the reference set of DGRs already collected from the literature, while the latter formed the main dataset analyzed in this study.

Successive rounds of the detection pipeline were used as follows. Candidate DGRs were first detected based on matches to Pfam Reverse-transcriptase domains (PF00078, PF07727, PF13456, and PF13655), with a score ≥20 for genomes and ≥30 for metagenomes^40^. Candidates with a RT sequence length of 250 to 550 amino acids and an imperfect repeat detected within 20kb of the RT in 5’ or 3’ (blastn hit ≥50bp with <99 % nucleotide identity), with one of the repeats within ≤1kb of the 5’ or 3’ end of the RT gene, were selected as putative DGRs. Sequences obtained from isolates were clustered with literature-derived references to remove redundancy (cd-hit^41^ v4.8.1, ≥95% amino acid identity). All candidate DGRs were included in a phylogenetic tree along with DGR and non-DGR references, based on a multiple alignment of RT amino acid sequences obtained with mafft^42^ v7.407 using iterative refinement (“einsi”), and built with FastTree^43^ v2 using the WAG substitution model. Since known DGRs formed a large monophyletic clade in this phylogeny, new sequences that branched outside of this clade and for which <75% of the mismatches between repeats were associated with A residues were considered as false positives and discarded. All other candidate DGRs were retained, and used to generate 6 new HMM profiles representing the main clades in the RT tree, using multiple alignment built with Muscle^44^ v3.8 and the hmmbuild tool from HMMER^37^ v3.2.1, with default parameters.

A second round of search was conducted on the same input dataset by using these new DGR RT HMM profiles instead of Pfam HMM profiles in the initial search (score ≥50 and e-value ≤1E-05). Putative DGRs were then selected as in the first round, except for the RT sequence length which was extended to range from 150 to 650 amino acids. After manual inspection and removal of false positive detections based on a phylogeny (as in the first round), another set of 4 HMM profiles were constructing. These new HMM profiles were used in a third and last round of search, which did not yield any new plausible candidate DGR, detecting only 7 previously undetected sequences that were all identified as likely false positives.

### Selection and annotation of reference genomes and metagenome sequences

For additional DGRs identified from IMG genomes, taxonomic classification was derived from the IMG taxonomy database. VirSorter^45^ v1.0.5 was used to identify which of these DGRs were encoded in proviruses: predictions of categories 1, 2, 4, and 5 were considered as viral, while predictions of categories 3 and 6 were provisionally listed as “putative viral” for the RT OTU-level aggregation (see below).

For DGRs detected on metagenome contigs, taxonomic classification of DGR-encoding contigs ≥3kb were derived from the automatic taxonomic annotation provided by IMG, based on majority ruling from gene-level best blast affiliations, i.e. each contig is affiliated based on individual gene affiliation up to the rank at which there is no majority affiliation anymore^31^. For contigs <3kb, taxonomic classification was set as “unclassified” because there are not enough predicted genes beyond the RT and DGR target on these contigs for a robust classification using the majority rule approach. Viral origin was predicted using a combination of VirSorter^45^ v1.0.5 (as for genomes), the Earth’s Virome pipeline^46^, and the inovirus detector pipeline^47^. All sequences identified as viral with the Earth’s Virome pipeline, the inovirus detector pipeline, or VirSorter categories 1, 2, 4, and 5 were considered “viral”, while sequences only predicted with VirSorter as categories 3 or 6 were listed as “putative viral” for the RT OTU-level aggregation (see below). For sequences not identified as viral by any pipeline, contigs ≥10kb were considered as “cellular”, while contigs <10kb were considered as “unknown”, based on previous benchmarks of viral sequence detection tools^45,46^. For contigs identified as viral, host taxonomic classification was determined as follows. For contigs ≥10kb, IMG blast-based taxonomy was considered as the predicted host taxonomy^48^. All sequences predicted as viral were also compared to IMG CRISPR spacer database^49^ using blastn^38^ with options -dust no and -word_size 7. Hits between viral sequences and CRISPR spacers with 0 or 1 mismatch over the entire spacer length were selected and used to infer host taxonomic classification of the corresponding viral sequence. In the rare cases where IMG and CRISPR matches taxonomic affiliations were inconsistent, IMG taxonomy was used. Viral contigs were also classified using vContact2^50^, to obtain a taxonomic affiliation of the virus itself (instead of the host) at the genus rank. vContact2 was run with “diamond” option to generate the PCs, clustering of VCs with cluster_one, and the reference database “ProkaryoticViralRefSeq94-Merged”, all other parameters left as default. Metagenome-derived DGRs were also associated to a biome type based on the original sample classification available in the Gold database^32^ (Supplementary Table 1).

In all analyses, the taxonomic classification used was the microbial one (i.e. “host” classification for viral contigs) at the domain and phylum ranks or equivalents. Members of the Candidatus Phyla Radiation (for bacteria) and DPANN (for archaea) were gathered in “Bacteria:CPR” and “Archaea:DPANN” groups (respectively) based on the supergroup classification proposed in ref. ^51^. Similarly, genomes classified as Bacteroidetes, Chlorobi, Cloacimonetes, Fibrobacteres, and Marinimicrobia were gathered in a “Bacteria:FCB” group, and genomes classified as Omnitrophica, Chlamydiae, Lentisphaerae, Planctomycetes, and Verrucomicrobia in a “Bacteria:PVC” group.

### Clustering and annotation of DGR based on RTs

The global DGR collection was clustered based on RT sequences, along with non-DGR RTs, representing a total of 33,342 RT sequences: 655 non-DGR RTs, 1,680 “reference” DGR RTs either from literature or from IMG genomes, and 31,007 from IMG metagenomes. These sequences were first clustered into “RT-OTUs” at 95% amino acid identity using cd-hit^41^ v4.8.1. Next, the representatives (longest sequences) from each RT-OTU were collected, and compared all-vs-all using blastp^38^ v2.9.0+. Blast hits with e-value <0.001 and with amino acid identity ≥50% (based on the whole query length) were used as input to an MCL clustering with inflation value 2.0 and amino acid identity percentage as edge weight^52^. The groups provided by MCL are designated as “RT-Clusters”.

RT-OTUs and RT-Clusters were associated with a genome type (viral or cellular), taxonomic classification, biome classification, and target gene (see below) as follows. Because DGRs were detected across >1,500 different bacterial, archaeal, and viral genera, and >90 different environment types, we opted to conduct global analyses at a coarse level, i.e. phylum-level for taxonomy and broad ecosystem types (e.g. “Aquatic:Freshwater”, “Host-associated:Rumen”) for biomes. Taxonomic classification of RT-OTUs was based on RT-OTU member affiliations using a majority rule, and an LCA if the top two affiliations had an identical number of members. A similar approach was used for the biome classification and the target PC, i.e. majority rule and LCA in case of a tie. For genome type, RT-OTUs with all members unknown were considered “unknown”, RT-OTUs with at least 1 “viral” or “putative viral” member and no cellular were considered “viral”, while others were considered “viral” or “cellular” based on a majority rule between viral and cellular members (if tied, the RT-OTU is considered as “unknown”). A consensus bias vector, representing the frequency of individual A, T, C, and G nucleotides in the TR for positions with mismatch, was also calculated by averaging the bias vector of RT-OTU members. For this average, we discarded cases in which the RT was found within 500bp of the contig edge or the target gene was found within 200bp of the contig edge, and the bias vector had an atypical A frequency <70%, as these likely represent misprediction of the TR/VR and/or target (based on manual inspection of these contigs). Similarly, in cases where RT-OTUs included both members with “typical” and “atypical” bias vectors, i.e. A frequency ≥70% and <70% respectively, the average vector for the RT-OTUs was calculated only with the “typical” ones. For RT-Clusters, a 2/3rd majority rule was applied based on the annotation of the RT-Cluster members. For multi-level data (taxonomy, biome, and target PC), the 2/3rd majority rule was applied first to the first level, then to the second level. In cases for which the majority value was found in less than 2/3rd of the RT-Cluster, the value was set as “unknown”.

### Enrichments of DGRs across taxa and biomes

Two different approaches were used to evaluate potential enrichment in DGRs of specific taxa and/or biomes. To link DGRs with specific taxa, the number of genomes affiliated to each taxon (at the same rank as for the DGR, see above) was estimated for each metagenome based on a list of 139 single-copy marker genes^53^. Briefly, for each metagenome, the total number of genomes for a taxon was estimated as the median number of single-copy marker genes affiliatied to this taxon, similar to the estimation performed in Anvi’o^54^. For each taxon, an enrichment in DGRs was calculated as the log2 ratio between the frequency at which DGRs were observed in this taxon (i.e. total number of DGR OTUs observed for this taxon divided by the total estimated number of genomes for this taxon across all metagenomes) and the average frequency of detection of DGRs across all taxa (i.e. total number of DGR OTUs divided by total estimated number of genomes across all metagenomes). The statistical significance of these differences in DGR frequency was evaluated using a Chi square test of independence (prop.test function in R on 2×2 contingency table for each taxon).

For biomes, the same set of single-copy marker genes were used to estimate a total number of microbial genomes in each metagenome. For each biome group (see above), a linear regression model was then fitted using the number of genomes as a predictor for the number of DGRs. For statistically significant fits (p-value <1E-4), an estimated number of DGRs per genome was derived based on the regression coefficient. Metagenomes with ≥40% of contigs ≥10kb identified as viral were excluded from this analysis as they likely derive from samples strongly enriched in viral genomes, for which the count of microbial genomes will not be reliable.

### Clustering and functional annotation of predicted target genes

For each DGR RT, target genes were identified by comparing the predicted VR repeat to CDS predictions available in the IMG database. If multiple VR repeats were detected for a single DGR RT, the one with the highest A mutation bias or, if tied, the closest to the DGR RT on the contig, was considered as the primary target. Genes associated with other VR repeats were then included as “secondary” targets if the VR repeat was associated with a plausible A mutation bias (≥75% of mismatches on A positions in the TR).

For *de novo* clustering of predicted targets, high-quality target sequences were first selected as genes longer than 300 nucleotides and not within 50bp of the edge of the contig, i.e. less likely to represent partial genes. These high-quality target genes were clustered at 99% AAI using cd-hit^41^ v4.8.1, then clustered in a two-step process as in ref. ^47^. Briefly, sequences are first clustered using MCL^52^ from an all-vs-all blastp, using blast score as edge weight and an inflation value of 2.0, then HMM profiles were built for these clusters and hhsearch^55^ v3.1.0 was used to identify similarities between clusters. This led to the definition of “superclusters” (i.e. clusters of clusters) based on a single-linkage clustering using similarities of ≥90% probability of ≥50% of the profile length or ≥99% over ≥20% of the profile length and 100 positions. Target sequences that were initially discarded because shorter than 300 nucleotides and/or within 50bp of the contig edge were then mapped to these superclusters using hmmsearch^37^ v3.2.1, with each sequence affiliated to the cluster with the highest score if ≥30.

Functional annotation of superclusters was obtained from analysis of the supercluster multiple sequence alignment and from the annotation of individual members. For the former, multiple sequence alignments were built using Muscle^44^ v3.8 after dereplicating the supercluster sequences at 90% AAI using cd-hit^41^ v4.8.1. These alignments were then used as input in hhsearch^55^ which compared the alignments to the Pdb70 v190918, Pfam v32, and scop70 v1.75 databases (database package downloaded in Feb. 2019 from the HH-Suite website). Each target sequence was also annotated the same way, using a direct hhblits^55^ comparison to the same Pdb70 v190918, Pfam v32, and scop70 v1.75 databases. In addition, individual target sequences were also searched for transmembrane domains and signal peptides using TMHMM^56^ v2.0c and SignalP^57^ v4.1, and compared to the Pfam v30 database using hmmsearch. Annotations were derived from hits with score ≥50 in hmmsearch or ≥90% probability in hhblits, except for hits overlapping the prediction VR region for which these cutoffs were lowered to ≥30 on score and ≥80% probability, in order to enable the identification of distantly related C-lectin folds. A target PC was considered as having a C-Lec fold VR if ≥5% of the PC members displayed a significant hit to a C-Lec fold reference overlapping with the predicted VR region. Similarly, a target PC was considered as functionally annotated outside of the VR if ≥5% of the PC members had a significant hit to a single reference sequence and/or domain.

A prediction of 3D structure was computed for selected target sequences using I-TASSER^58^ 5.1, using default reference libraries and 25 hours-long simulations. The average conservation of residues in target clusters was based on the multiple alignments generated for cluster annotation (see above). For comparing conservation within and outside of the VR regions, a predicted “extended” average VR region was defined by adding 200 residues upstream and 20 residues downstream of VR regions predicted on individual sequences. The size of this “extended” VR regions was defined based on the coordinates of predicted VRs and surrounding conserved C-Lec fold on reference DGRs.

The association between RT, TR-VR, and target protein sequences was evaluated as follows. Predicted TR sequences associated with high-quality targets and typical mutational bias (see above) were compared using all-vs-all blastn^38^ v2.9.0+ with options adapted for short sequences (“--dust no -- word_size 7”). The global nucleotide identity between two TR sequences was then calculated based on the number of identical residues in the best blast hit compared to the length of the shortest TR sequence of the pair.

### DGRs identified in genome bins

When available, automatically-generated genome bins were searched for DGR-encoding contigs. Briefly, genome bins were automatically generated for public metagenomes on IMG, using metabat^59^ v0.32.4 for binning with a 3,000 bp minimum contig cutoff, contig coverage information, and parameter ‘-superspecific’ for maximum specificity, checkM^60^ v1 for quality estimation, and gtdb-tk^61^ v0.3 for taxonomic assignment. Only medium- and high-quality bins according to the MIMAG^62^ criteria were included. DGRs encoded on contigs identified as entirely viral were not considered in this process, since previous studies have indicated that these contigs are often binned incorrectly. Overall, 13,180 MAGs were searched, and 1,509 were found to include at least 1 DGR locus. For metagenomes including at least 1 genome bin with a DGR-encoding contig and at least 10 genome bins, the relative abundance rank of each genome bin was determined as follows: for all MQ and HQ genome bins identified in the metagenome, the bin coverage was estimated as the median coverage of all contigs. The genome bins were then ordered based on this median coverage of contigs to determine the rank(s) of genome bin(s) encoding DGRs.

The diversity of DGR-encoding genomes identified in human gut samples was evaluated through an RNA polymerase B (RpoB) tree. RpoB protein sequences were first identified in isolate genomes and genome bins associated with human gut and encoding a DGR of clade 1, 4, or 6, based on significant hits to the pfam domain PF04563 (hmmsearch score ≥50). A multiple alignment was then built with MAFFT^42^ v7.407 using default parameters, automatically trimmed using TrimAl^63^ v1.4.rev15 with the --gappyout option, and used as input to build a tree with IQ-Tree^64^ v1.5.5 with built-in model selection (optimal model suggested: LG+R6).

The gene content of DGR-encoding genomes was evaluated based on metagenome bins as follows. For each metagenome including at least one Clade 5 DGR in a genome bin, the number of proteins affiliated to each COG in each MQ or HQ bin was tallied. The number of proteins associated with each COG category (level 1) was then compared between DGR-encoding genome bins and non-DGR-encoding genome bins for each metagenome using a Kolmogorov-Smirnov test and Cohen’s effect size.

### Phylogenetic analyses

For RT phylogeny, a representative of each RT cluster was selected as the sequence with the highest score when compared to the RT cluster hmm profile, or the longest sequence in case of ties, first among the references if available, then among the new sequences if no reference was present in the RT cluster. An RT tree was then built with IQ-Tree^64^ v1.5.5 using the built-in model selection (optimal model suggested: VT+F+R10), based on an amino acid multiple alignment computed with MAFFT^42^ v7.407 using the einsi mode, and automatically trimmed using TrimAl^63^ v1.4.rev15 with the --gappyout option. Sequences from the non-DGR RT reference set (see above) were included in the alignment, with the exception of sequences identified as “unknown” or “unclassified” by Wu et al., as these led to a lower quality alignment and long branch attraction issues in the resulting tree.

The distribution of DGR features (genome type, taxonomy, target, and biome) across the tree was analyzed by computing unweighted Unifrac distances^65^ between all pairs of values for each features and comparing these with the distance for the same pair of values on 100 randomly shuffled trees. Ancestral state reconstructions were conducted using the R package phytools^66^ v0.6-99 with the options model=“ER” and type=“discrete”, separately for each feature.

A similar pipeline was used to build the trees of target sequences from PC_00003. Briefly, for PC_00003 targets, high-quality target protein sequences (see above) clustered into PC_00003 were gathered and dereplicated at 80% amino acid identify using cd-hit^41^ v4.8.1. A multiple alignment of representative sequences was then generated with MAFFT^42^ v7.407 using the einsi mode, automatically trimmed using TrimAl^63^ v1.4.rev15 with the --gappyout option, and used as input to build a tree with IQ-Tree^64^ v1.5.5 with built-in model selection (optimal model suggested: WAG+F+R10).

### Association between DGR presence or features and genome characteristics

To evaluate potential links between the presence of a DGR and the original biome or taxonomic classification of a genome, a random forest classifier was trained to predict from a genome’s features whether or not it encodes a DGR. A training set for the model was built based on DGR and single-copy marker genes observed in metagenomes as follows: first a metagenome was selected at random, then a genome taxonomy was selected at random with probability weighted based on the affiliation of single-copy marker genes in this metagenome (see “Enrichments of DGRs across taxa and biomes”), and this genome was considered to encode a DGR or not based on the overall frequency of DGRs detected in genomes of the same taxonomy in this metagenome. This process was repeated until a dataset of 1,000 DGR-encoding and 1,000 non-DGR-encoding genomes was obtained. This allowed for the generation of a balanced dataset which still reflected actual frequency of DGRs detected across taxa and metagenomes. Random forest classifiers were generated with the randomForest^67^ R package v4.6-14 using as predicting features a combination of taxonomy and/or biome features, either at 1-level (e.g. “Aquatic” or “Bacteria”) or 2-level (e.g. “Aquatic:Freshwater” or “Bacteria:Proteobacteria”) classes. All classifiers were built using 500 trees with all other options default. The Out-of-bag confusion matrix automatically generated was then used to evaluate the models’ prediction accuracy.

### Diversity estimation of DGR target loci

Nucleotide and amino acid diversity evaluation was conducted on the metagenome-derived DGRs with contig length ≥10kb and median coverage ≥20x. The coverage cutoff was applied to ensure that single nucleotide polymorphisms (SNVs) could be called with enough certainty, while the length cutoff was used to ensure that enough surrounding genes were available to evaluate background microdiversity for the DGR-encoding genome. For this analysis, combined assemblies (i.e. datasets obtained by combining reads from multiple samples), metatranscriptomes, and viral metagenomes were not included. The final set included 6,901 DGRs, with representation of all DGR clades (1,968 DGRs from DGR_Clade_1, 972 from DGR_Clade_2, 1,359 from DGR_Clade_3, 1,147 from DGR_Clade_4, 1,095 from DGR_Clade_5, and 350 from DGR_Clade_6).

For DGRs found on contigs ≥20kb, a region of 20kb around the RT (i.e. up to 10kb in 5’ and 3’) was extracted and used in these analyses. Reads from the original metagenome were first recruited to the contig (or selected contig subset) using bwa^68^ v0.7.17-r1188 (default parameters). Reads which matched the reference sequence on at least 50% of their length were then extracted using filterBam (https://github.com/nextgenusfs/augustus/tree/master/auxprogs/filterBam) and realigned against the same reference sequence using bbmap^69^ v38.73 to obtain a global alignment of the read to the reference contig instead of the local/soft-clipped alignment provided by bwa (bbmap options “vslow minid=0 indelfilter=2 inslenfilter=3 dellenfilter=3”, see Supplementary Text). This global alignment was required to accurately estimate SNVs in regions with a high number of mismatches, such as VRs with many different variants in the population. Typically, in these regions, local alignment tools will either trim the mapping to remove these mismatch-containing regions, or “soft-clip” them, i.e. mask them in the resulting sam file, which eventually means that no SNV will be called in these variable regions. Instead, by re-aligning the same reads with a global alignment algorithm, all positions from the read will be considered and SNVs can be identified.

SNVs were called using bcftools^70^ v1.9 ‘mpileup’ and ‘call’ functions (options “-A -Q 15 -L 8000 -d 8000” for mpileup, “--ploidy 1 -A -m” for call). Only SNVs for which the alternative allele was supported by ≥4 reads or ≥1% of the reads (whichever was smaller) were further considered. These SNVs were then classified as synonymous or non-synonymous based on the available gene prediction, and used to calculate pN/pS for each gene as in Schloissnig et al.^71^. Another set of SNVs was called using FreeBayes^72^ v1.3.1 using the options “--ploidy 1 --min-base-quality 15 --haplotype-length 0 -- min-alternate-count 1 --min-alternate-fraction 0 --pooled-continuous --limit-coverage 8000” and the same cutoff on read representation. The two SNV sets were found to be mostly overlapping (see Supplementary Text), and the bcftools SNVs were used in all subsequent analyses.

To evaluate DGR activity, an enrichment of the VR locus in non-synonymous SNVs was calculated as follows. For each DGR, the frequency of non-synonymous SNVs (i.e. average number of non-synonymous SNVs per position) observed across all genes for the contig (or contig subset) was first calculated. A poisson law was then used to compare the number of non-synonymous SNVs observed in the VR locus to an expected number of non-synonymous SNVs based on surrounding genes. All cases for which the number of non-synonymous SNVs observed in the VR locus was significantly higher than expected by chance (poisson law probability <0.05) were considered as observations of DGR activity. For cases where different metagenomes were available from the same sample, only the observation with the highest coverage was selected and used in DGR activity evaluation.

To further explore the dynamics of VR loci in nature, 130 datasets were identified as longitudinal sampling that enabled tracking of DGRs across time in the same system (Table S7). For these, a similar mapping approach was used as described above, but including all metagenomes from a single subject (for human-associated metagenomes) or location (e.g. geographic coordinates and water layer). The same approach based on non-synonymous SNV frequencies was used to evaluate DGR activity for each individual sample, except that the minimum median coverage was set at 10x. For cases in which multiple metagenomes were available for a single subject/location and time point, the one with the highest coverage was used for each DGR. In addition, nucleotide diversity (Π) was calculated for each VR locus as the average nucleotide diversity observed for each position. To evaluate changes in the VR locus between samples, amino acid variants were called using Anvi’o^54^ v6.1 based on the same read mappings. This enabled us to accurately evaluate the frequency of individual amino acid residues, including the ones for which multiple positions in the codon were variable.

### Data processing and visualization

Plots and charts were generated in R^73^ v3.6.1 using the ggplot2^74^ v3.2.1 package, while phylogenies were visualized using iToL^75^ v4 (https://itol.embl.de/) and predicted protein structures were visualized with UCSF Chimera^76^ v1.11.2. Several programs used in this study benefited from the GNU parallel tool^77^. The genome maps from Supplementary Fig. S4 were generated using Easyfig^78^ v2.2.3.

### Data and scripts availability

Scripts used in this study are available at https://bitbucket.org/srouxjgi/dgr_scripts/. All metagenome assemblies are available through IMG (https://img.jgi.doe.gov) using accession numbers listed in Supplementary Table S1. Additional supplementary datasets include:

All_RTs.faa: fasta file of amino acid sequences for all RTs used in the analysis (DGR and non DGR) All_DGR_targets.faa: fasta file of amino acid sequences for all predicted DGR targets. Note that this file includes all sequences predicted by the automatic pipeline to be a DGR target, before any of the manual curation step mentioned in the Methods section All_DGR_TR_VR.fna: fasta file of nucleotide sequences for TR/VR pairs. As for the targets, this file includes all sequences predicted by the automatic pipeline, before any of the manual curation steps mentioned in the Methods section and Supplementary Text.

The same files are also available at: https://bitbucket.org/srouxjgi/dgr_scripts/src/master/Companion_datasets/.

## Acknowledgments

We thank Dr. Sean Crowe, Dr. Laura A. Hug, Dr. Kelly C. Wrighton, and Prof. Katherine D. McMahon for providing detailed information on unpublished metagenomes. The work conducted by the U.S. Department of Energy Joint Genome Institute is supported by the Office of Science of the U.S. Department of Energy under contract no. DE-AC02-05CH11231. This research used resources of the National Energy Research Scientific Computing Center (NERSC), a U.S. Department of Energy Office of Science User Facility operated under Contract No. DE-AC02-05CH11231. B.G.P. was supported by the Marine Biological Laboratory, by the National Science Foundation’s XSEDE computing resource (award DEB170007), and through a Challenge Grant from the California NanoSystems Institute at the University of California Santa Barbara. M.A.O. acknowledges funding support from the National Science Foundation (NSF) (MCB-1553721), the Camille Dreyfus Teacher-Scholar Awards Program, and the California NanoSystems Institute (CNSI) Challenge Grant Program, supported by the University of California, Santa Barbara and the University of California, Office of the President. This work was part of the DOE Joint BioEnergy Institute (http://www.jbei.org) supported by the Office of Biological and Environmental Research of the DOE Office of Science through contract DE-AC02– 05CH11231 between Lawrence Berkeley National Laboratory and the DOE. R. C. and M. A. A. work was supported by the Australian Research Council (DP150100244) and the Australian Antarctic Science program (project 4031). The work in the Hallam Lab was performed under the auspices of the US Department of Energy (DOE) Joint Genome Institute, an Office of Science User Facility, supported by the Office of Science of the U.S. Department of Energy under Contract DE-AC02-05CH11231 through the Community Science Program (CSP), the G. Unger Vetlesen and Ambrose Monell Foundations, the Natural Sciences and Engineering Research Council of Canada, Genome British Columbia, Genome Canada, and Compute Canada and the Canada Foundation for Innovation through grants awarded to S.J.H.

## Supplementary Text

### Iterative detection of DGR in IMG public genomes and metagenomes

Overall, up to 3 rounds of DGR detection were performed for both genomes and metagenomes. The first round of detection was based on known HMM profiles of DGR RTs, while after each round, new profiles were generated from the DGR RT sequences collected, and used as references for the next searches (see Methods). For whole genome shotgun sequences (i.e. “IMG isolates”), the first round of searches identified 2,793 candidate DGR sequences, the second 442 candidates, while the third yielded only one candidate that was identified as a false-positive, hence no further searches were performed. For metagenomes, the first round identified 45,704 candidates, the second 10,964 candidates, while the third round provided seven candidates which were all identified as false-positives. Overall, DGRs were detected across 1,129 genomes and 2,684 metagenomes.

For both genomes and metagenomes, false-positive detections were mostly associated with RTs encoded on eukaryote genomes, especially in regions containing multiple imperfect repeats with seemingly random mismatches and both repeats within predicted CDS, as opposed to the A bias and one intergenic repeat of typical DGRs. When included in a phylogenetic tree based on RT protein sequences, these candidates formed a clade outside of all known RTs, and in particular branched outside of the known DGR clade. Because these sequences are likely not representing genuine DGRs despite the presence of nearby repeats, the choice was made to only retain sequences with a typical DGR mismatch profile (i.e. enriched in A mismatches) or branching within the known DGR clade, while all other candidates were excluded.

### Taxonomic distribution of DGRs detected from metagenomes

While a majority of DGR-encoding contigs could only be affiliated to the phylum or class rank, a total of 4,755 were affiliated up to the genus rank, distributed across 369 bacterial genera and 9 archaeal genera. Even though 48% of these affiliations were to only 3 genera (*Bacteroides, Pseudomonas*, and *Prevotella*), both because of a high prevalence of DGRs in these taxa and an over-representation of these sequences in the metagenomes mined, the less common genera revealed new DGR-encoding taxa (Supplementary Table 2). These included notably members of the *Fibrobacter* genus, key actors in the degradation of cellulose compounds in ruminant animals, for which DGRs have not yet been described and explored. This dataset also included additional examples of DGR for ecologically-relevant genera within the phyla *Chlorobi* (e.g. *Pelodictyon, Chlorobaculum*, and *Chlorobium*), *Actinobacteria* (e.g. *Bifidobacterium, Colinsella*, and *Gardnerella*), and *Nitrospirae* (e.g. Candidatus *Magnetobacterium*), for which only a handful of examples have been reported. Within the Archaea domain, DGRs were associated with various members of the *Euryarchaeota* phylum and DPANN supergroup.

For this taxonomic affiliation, viral-encoded DGRs were associated with the taxon of their host when available, based on the detection of an integrated provirus or matches to known CRISPR spacers (see Methods). However, the viral genomes encoding these DGRs can also be classified in a separate viral taxonomic framework. When affiliated, nearly all DGR-encoding viruses (n=3,218) were classified in the *Caudovirales* order (either affiliated to an existing *Caudovirales* genus or connected to the main *Caudovirales* component in vContact2 network). Notably, while many giant viruses have been recently identified from metagenome assemblies^79^, no DGR were detected in these genomes or co-localized with an NCLDV marker gene, suggesting that DGRs are rare or absent from these large eukaryotic viruses despite their ability to exchange genes with bacteriophages. The only exceptions were two sequences(Meta_3300029305_Ga0307249_1003718615and Meta_3300018430_Ga0187902_100048955) identified as putative inoviruses because of the presence of an inovirus ATPase marker nearby^47^. However, in both cases, the metagenome contig was too short to distinguish whether the DGR was encoded by the inovirus genome or by a neighboring *Caudovirales* prophage, as previously observed^47^. Hence, overall, the available affiliations of DGR-encoding virus contigs suggest that these elements are mostly, and maybe exclusively, encoded by *Caudovirales*.

### Homogeneity of DGR OTUs and Clusters based on RT sequences

DGR OTUs were defined based on a 95% AAI clustering of the RT sequences, and were homogeneous in terms of taxon, biome, and genome type (i.e. viral vs cellular). Specifically, 99%, 91%, and 94% of non-singleton DGR OTUs were associated with a single taxon, biome, and genome type, respectively (Supplementary Fig. S2). A majority (60%) of DGR RT sequences remained as singletons after this 95% AAI clustering, and >88% of DGR RTs were found in OTUs comprising less than 5 sequences, illustrating how large the DGR RT sequence space is.

A similar pattern was observed for DGR clusters (≥50% AAI groups), of which 99%, 85%, and 96% were associated with a consistent taxon, biome, and genome type, respectively (Supplementary Fig. S2). The lower percentage of consistency for the biome feature was due in part to overlap between connected environments, e.g. “Landfill” and “Groundwater”, as well as the detection of DGR clusters with a broad ecological distribution such as Meta_3300009658_Ga0116188_10269693, which includes members from wastewater treatment plants, biogas reactors, saline and freshwater lakes, elephant gut microbiome and moose rumen microbiome (Supplementary Table 4). While an exception, this suggests that at least some DGR clusters are broadly distributed in the environment. Conversely to the OTU clustering, most DGR RTs were found in a cluster comprising 2 or more sequences, and only 52% of DGRs were found in clusters including less than 5 sequences. In addition, the 11 largest clusters alone gathered >36% of all sequences, illustrating how this dataset enabled the detection of the predominant groups of DGRs in the environment. Consistently, the 11 largest clusters all included at least one reference sequence (Supplementary Fig. 1).

### Definition of major DGR clades

In order to partition the large RT phylogeny into meaningful clades, we sought to leverage the key features of each DGR cluster including genome type, taxon, and biome. We first mapped each parameter to the RT phylogeny rooted on non-DGR RTs and verified that the distribution of each parameter across the tree was statistically structured and not random (non-weighted Unifrac p-value <1E-03). Then, we reconstructed ancestral states for each parameter at each node throughout the tree, and identified deep-branching clades with ancestral states predicted with ≥95% confidence as the main groups of DGRs. Taken together, the different features mapped onto the tree suggest the successive emergence of 6 major DGR clades from a single origin (Fig. 1A & B).

Based on these ancestral state reconstructions, the deepest-branching clade of DGRs (DGR clade 1) was found primarily in host-associated microbiomes (>99% confidence) and encoded on viral genomes (>99% confidence) infecting mostly *Bacteroides* and *Firmicutes* hosts. DGR clade 4 showed similar characteristics with a clear association with host-associated microbiomes (>99% confidence) and *Firmicutes* (>99% confidence), and most sequences found in a viral genome. Two large groups of cellular-encoded DGRs (DGR clades 2/3 and DGR clade 5) branched next to DGR clade 4 suggesting two potentially distinct horizontal DGR transfer events from viruses to bacteria/archaea. In the group including clades 2 and 3, the deep-branching nodes are associated with aquatic (>99% confidence) cellular-encoded DGRs (>99% confidence) affiliated to the CPR (Candidate Phyla Radiation) taxon (>96% confidence). However, another type of DGR is nested within these CPR-encoded DGRs, also originally associated with aquatic environments (>99% confidence) but affiliated to other bacteria taxa, especially *Proteobacteria*. The deep-branching CPR-associated DGRs were identified as “DGR Clade 2”, while the latter clade was identified as “DGR Clade 3”. The other group of cellular-encoded DGRs, “DGR clade 5” branched separately from clades 2 and 3 but was also predicted to originally represent aquatic (>99% confidence) cellular-encoded (>98% confidence) DGRs. A single clade of viral-encoded (>99% confidence) DGRs branched within it however, suggesting a secondary transfer this time from cellular to viral genomes. This latter clade was identified as “DGR Clade 6”.

### Distribution of reference DGRs across newly defined clades

While the diversity of DGR RTs was vastly expanded by metagenome-derived sequences, each of the 6 DGR clades included at least one reference and/or laboratory-characterized DGR sequence^2^. The original DGR identified in Bordetella virus BPP1^3^ was affiliated in DGR Clade 6, along with other reference DGRs mostly identified in prophages from *Proteobacteria* and *Firmicutes* genomes, targeting viral structural proteins (Target PC_0002 and PC_00010, Fig. 2, Supplementary Fig. 8). The *Treponema* DGR^9^ was affiliated to DGR Clade 5, along with DGR sequences identified in *Spirochaetes, Cyanobacteria*, as well as *Archaea* and associated archaeoviruses^5^. Consistent with the majority of DGR Clade 5 sequences, these reference sequences represented the main clade of cellular-encoded DGRs associated with membrane-bound target proteins (Target PC_00001). DGR Clade 4 included another set of DGRs previously identified on prophages, mostly from *Firmicutes* and *Actinobacteria*, and targeting viral structure proteins. The *Legionella* DGR^8^ was affiliated in DGR clade 3, along with other references from *Proteobacteria* and *Bacteroidetes* targeting uncharacterized proteins gathered in Target PC_00009. Interestingly, a number of these references correspond to prophage-encoded DGRs, while others, such as the *Legionella* DGR, are encoded on regions of the cellular chromosome that are predicted as “putative mobile genetic element”. This suggests that this DGR may have been frequently exchanged between cellular and viral genomes, or may have originated from a provirus and retained on cellular genomes while the rest of the provirus decayed. DGR Clade 2 gathered sequences identified in the CPR group^6^ and associated with various uncharacterized targets including Targets PC_00019, PC_00020, PC_00024, PC_00027, PC_00029, and PC_00040. Finally, DGR clade 1 included references from *Bacteroidetes, Firmicutes*, and *Actinobacteria* prophages, targeting mostly viral structural proteins (Target PC_00002, Fig. 2, Supplementary Fig. 8).

### Distinguishing prediction errors from genuine atypical TR-VR sequences

While our detection approach did not rely on the typical mutational bias of DGRs, i.e. mutations associated with A residues, nearly all (99%) DGR clusters displayed a strong bias towards A mismatches (Fig. 1B). The few TR-VR pairs which did not show this pattern could arise either from different mispredictions of the TR-VR regions or from a genuine DGR not constrained by the typical A mutation bias. Upon manual inspection, different scenarios leading to mispredicted TR-VRs were identified:

- Errors in the predicted CDS in the VR region can lead to an incorrect “T” bias, i.e. most of the mismatches correspond to “T” in the TR because the putative target gene is predicted on the opposite strand of the real target gene. This is especially common for VR regions situated near the edge of a contig (i.e. within the first or last ∼ 200bp), and we thus discarded such mutation bias profiles based on partial target genes predicted on the edge of a contig.
- In other cases, the CDS prediction overlapping the VR region seemed correct, however it is unlikely to be a genuine DGR target based on functional annotation and a lack of similarity to any other DGR target (known or predicted). In this case, it is most likely that the TR-VR pair was wrongly identified, and the associated mutations biases were also excluded.
- For TR-VR with a plausible target gene and not on the edge of a contig, several cases were identified for which the blast hit used to define the TR-VR region likely extended past these regions. While the TR-VR initially identified did not display an A bias, an internal subset of the alignment could be identified upon manual inspection with (near-)exclusively A mismatches. Most likely in these cases, near-identical regions next to the TR-VR led to this misprediction, and the bias was corrected to reflect the one of the “inner” TR-VR.
- Finally, some TR-VR pairs were associated with a plausible target gene and did not include any subset with an A bias. No other mutation bias was apparent in any of these TR-VR pairs, i.e. the mismatches observed did not show any enrichment in a specific nucleotide either in the TR or the VR sequence. These could represent either TR-VR sequences that accumulated mutations beyond the ones introduced by the DGR RT, or DGR RTs for which the incorporation of random nucleotides is not strictly associated with A residues. The small numbers of such “atypical” DGRs and their sparse distribution across the tree indicates however that the tendency of DGR RTs to incorporate random nucleotide specifically at template A-residues is both ancestral and conserved, thus most likely associated with biochemcial and/or structural constraints of the RT enzyme itself^11^.

### Target clustering and annotation

To avoid over-estimating the diversity of target proteins, we first restricted the analysis to only “high-quality” targets, i.e. predicted target genes longer than 300 nucleotides and not within 50bp of the edge of the contig. This selection was designed to limit the inclusion of incorrect targets due to partial and/or mispredicted CDS in metagenome contigs. Dereplication (99% AAI) of these high-quality targets led to a dataset of 15,559 non-redundant targets used as input in the two-step clustering process (see Methods). After protein clustering, most (92.18%) high-quality targets clustered in 1 of the 24 largest PCs, which were all associated with plausible functional annotation for DGR targets, and thus further considered as likely genuine DGR targets. Other target sequences found in smaller PCs and/or singletons could be either genuinely rare types of targets or cases for which the target CDS or TR-VR regions are misidentified. Because these sequences are mostly originating from short contigs for which genes and DGR features prediction are challenging, we opted to consider these targets as “Rare” and not analyze them further.

To evaluate the expansion of DGR target space obtained here, we compared the functional annotation obtained on the 24 largest target PCs with the functions of DGR targets previously reported^2^. In addition to conserved domains already detected on reference sequences, 23 domains were newly identified on >5 predicted target sequences. These included conserved domains within S-Layer-containing proteins (PDB 4QVS_A), PEGA domains (Pfam PF08308), putative bacterial lipoproteins (Pfam PF05643), putative glutamyl endopeptidases (PDB 1WCZ_A), serine proteases (PDB 3STI_A and 6BQM_A), and other types of viral structural proteins (PDB 4V96_AG, Pfam PF03906). Overall these confirm that DGR targets are extremely diverse, but are mainly associated with carbohydrate-binding proteins embedded in virions and cell membranes.

### Examination of putative Ig-like VR domains

Wu et al.^2^ previously reported two ‘categories” of VRs which were predicted to adopt an Immunoglobulin-like (Ig-like) fold (Ig1 and Ig2). These included respectively 36 and 9 non-redundant targets, with 3 types of domain organization: 25 displayed a C-terminal VR within a short (≤250 aa) protein, 4 included a C-terminal VR with a long (>250 aa) uncharacterized N-terminal part, and 17 had a N-terminal VR followed by a long (>250aa) uncharacterized C-terminal region^2^. As typical for DGR targets, despite the similarities observed between the VR regions, most of these sequences could not be annotated. Wu et al. however noted that a structure prediction with Phyre 2 suggested that some of these VRs (3 Ig1 and 5 Ig2) may overlap with an Ig-like domain.

In our analysis, Ig1 and Ig2 targets were found in two different PCs, PC_00003 (24 Ig1, 8 Ig2) and PC_00008 (12 Ig1), with one exception (one Ig2 target was clustered in PC_00062, considered a “rare” target in our analysis). Both PC_00003 and PC_00008 are clearly associated with viral-encoded DGRs, which is consistent with previous analysis of Ig1 and Ig2 VRs^2^. However, with PC_00003 and PC_00008 including 1,770 and 416 non-redundant high-quality sequences respectively, this extended dataset provided a unique opportunity to further understand the putative link between VRs and Ig-like domains.

When annotating these targets using hhblits, members of PC_00003 and PC_00008 did not display any conserved domain overlapping the VR region (Supplementary Table 5). Both members of PC_00003 and PC_00008 had hits to functional domains related to carbohydrate binding outside of the VR regions however, including several with Ig-like domains (Supplementary Table S5). Several types of phage tail fiber proteins have been previously shown to harbor similar Ig-like fold domains^80^, which is consistent with expected function of DGR targets. Specifically, this type of tail fiber was shown to vary in length, especially through the addition and removal of one or several Ig-like domains in the C-terminal region of the protein.

Accordingly, when building a phylogeny of target sequences from PC_00003 and mapping the domain organization of these targets to the tree, two large clades of sequences longer than average and typically including one or several Ig-like domains immediately downstream to the conserved VR domain can be observed (Supplementary Fig. S7). This would be consistent with the original target displaying a typical C-terminal VR region, and progressively increasing in length through the downstream addition of additional carbohydrate-binding domains.

We wondered however whether Ig-like domains immediately next to VR regions could lead to erroneous structure predictions including an incorrect overlap between VR and Ig-like domains. When we repeated the structural prediction of Ig1 and Ig2 proteins using Phyre2, we observed that all cases for which an Ig-like fold was predicted as overlapping a VR region corresponded to sequences for which multiple Ig-like domains were identified (both by hhblits and Phyre2) downstream from the VR, and found in one of the two clades of sequences longer than average (Supplementary Fig. S7). The structures obtained were also of relatively low quality (as also noted by Wu et al.^2^), and based on templates often consisting of multiple successive Ig-like domains. Conversely, for all Ig1 and Ig2 sequences with a C-terminal VR or with a VR without nearby Ig-like domain, no prediction of an Ig-like fold was obtained. These Ig1/Ig2 sequences are thus most likely phage tail fibers with a non-Ig-like conserved VR domain for which no characterized fold can be identified.

### Analysis of DGR-encoding metagenome bins

Two analyses were conducted based on DGR identified in IMG genomes bins (see Methods). First, genome bins were used to evaluate whether DGR-encoding viruses infected dominant and/or rare genomes in human gut samples. A total of 124 human gut metagenomes were selected which included at least 10 MQ/HQ bins, at least 1 DGR-encoding bin, and for which coverage information was available. While these metagenomes included 10 to 37 bins, DGR-encoding bins were frequently among the most abundant genomes observed. Specifically, DGRs were identified in one of the 3 most abundant bins for 68 of these 124 metagenomes (55%). While this pattern may be potentially biased by the fact that genome bins with higher coverage may have a higher completeness than bins with lower coverage, importantly, it was not observed across other biomes. Specifically, when evaluating the number of metagenomes for which a DGR was identified in one of the 3 most abundant bins, these only represented 17 of the 66 qualified metagenomes (26%) from other host-associated samples (i.e. non-human gut), 44 of the 139 qualified metagenomes (32%) from aquatic samples, and 17 of the 67 qualified metagenomes (25%) from engineered samples, compared to 55% for human gut samples. Hence, this pattern is likely not associated with a systematic genome binning bias, and DGRs seem to be specifically associated with abundant members of the community in human gut microbiomes.

Next, we used genome bins to evaluate differences in gene content between DGR-encoding and non-DGR-encoding genomes for Clade 5 DGRs, which are primarily identified in aquatic biomes. Compared to MQ/HQ genome bins from the same metagenomes that do not encode a DGR, genome bins encoding a Clade 5 DGRs displayed a significant enrichment in key COG categories associated with copiotrophic and/or particle-associated lifestyle. Specifically, genome bins which included a Clade 5 DGR showed a higher percentage of genes assigned to COG category N “Cell motility”, T “Signal transduction mechanisms”, and V “Defense mechanisms” compared to other bins from the same metagenomes (ks-test p-value ≤1E-5, cohen’s effect size ≥0.2), all categories previously highlighted as enriched in copiotrophs^81^. Importantly, the same pattern, i.e. significant enrichment in COG categories N, T, and V, was observed for bins including the other main clade of cellular-encoded DGRs, DGR clade 3, but not for any other clade. This indicates that these patterns are not a systematic bias of genome binning, but instead reflect key features in terms of gene content of micro-organisms encoding DGRs of clade 3 and 5. Given that the target genes of DGR clade 5 are typically membrane-bound, it is tempting to speculate that at least some of the DGRs in these clades drive hyperdiversification of surface protein directly involved in particle binding, and would thus broaden the range of particles and/ or other microbial cells to which a microbe could bind.

### Evaluating DGR activity from read mapping

Because of the high number of SNVs concentrated in a short region, mapping pipelines for VR sequences must use specific parameters to allow more mismatches and avoid under-recruiting short reads to the hypervariable reference. To achieve this, we used here a two-step mapping process. First, all reads were mapped using bwa to recruit all reads matching, even partially, to the reference sequence (i.e. based on local alignment). Then, reads which were locally aligned on at least 50% of their length were mapped against the same reference using bbmap with parameters tuned toward optimization of the read global alignment and tolerating mismatches (see Methods). We verified whether most reads from VR regions were likely recovered by comparing the read depth of VR regions to the one of surrounding genes from the same contig (Supplementary Fig. S9). Since some metagenomes will display variable read coverage along a single contig, e.g. due to PCR amplification of the library^82^, we established a 95% confidence interval for coverage along each contig based on the average coverage of non-target genes minus 2 standard deviations, and considered as “low coverage” cases in which the VR coverage was below this cutoff. Pragmatically, we considered VR region with a coverage below the average minus 2 standard deviations cutoff as unexpectedly low, and likely reflecting an incomplete recruitment of VR reads. Overall, >91% of VR regions displayed a coverage above the 95% confidence interval lower bound, confirming that most reads coming from these VR regions had been recovered.

To further verify that SNVs could be robustly called in VR regions, we compared the number of SNVs detected by two different tools: bcftools mpileup/call and freebayes (see Methods). Overall, both SNV calling approaches produced very similar result (Supplementary Fig. S9B), resulting in a Pearson correlation coefficient of 0.873 (95% confidence interval: 0.868-0.879, p-value <2.2e-16) between SNV densities for individual VR regions. We thus proceeded using only one of these SNV sets, and opted to use the more conservative one given the parameters used here, i.e. bcftools (Supplementary Fig. S9B).

In order to measure the selective constraints exerted on individual genes and/or VR regions, we first relied on the known pN/pS metric, calculated as in Schloissnig et al.^71^. For non-target genes, this pN/pS ratio was on average 0.16 (95^th^ percentile=0.60), as expected for microbial genes evolving under long-term purifying selection. By contrast, pN/pS ratio average was 2.83 for target genes. Notably, pN/pS calculations were sometimes impossible to calculate because of an absence of synonymous SNVs. While in the case of non-target genes, the absence of synonymous SNVs was mostly (>90% of the time) associated with an absence of non-synonymous SNVs, this was not the case for VR regions, of which 51% displayed ≥1 non-synonymous SNVs but 0 synonymous SNVs. In order to include these sequences in the activity estimation, we opted to use an “enrichment in non-synonymous SNVs” statistics comparing the density of non-synonymous SNVs in a VR region to the one in surrounding non-target genes. The two approaches were largely congruent, as for cases in which both could be calculated, the 296 sequences without an enrichment in non-synonymous SNVs all had low pN/pS (median=0.39), while the 2,417 sequence with an enrichment in non-synonymous SNVs had a high pN/ pS (median=2.48).

Finally, we searched for time-series datasets that could provide insights into the population diversity of DGR loci through time. We first identified time series among our metagenome set, and linked each sample to its “subject/location”, i.e. individual patient for human cohorts, individual bioreactor for laboratory incubations, individual water body and/or depth layer for lakes (Supplementary Table 6). Individual water layers were used as subject/location when they represented distinct ecological conditions, or were grouped as a single subject/location otherwise to avoid duplicate observations. Candidate DGRs for time series analysis were identified based on DGR OTUs which included members assembled from multiple datasets of a single subject/location. For these, reads from all datasets associated with the subject/location were mapped to the same DGR OTU representative sequence, and the longitudinal analysis was conducted if the median coverage of this sequence was ≥10x in ≥2 time points. Overall, 563 DGR OTUs were analyzed this way across 130 time series, and covered all DGR clades (the lowest number of DGR OTUs was for DGR clade 2, with 47 OTUs).

### Definition of activity categories for time series

The activity of DGR analyzed as part of a time series were evaluated based on single amino acid variants called using Anvi’o (see Methods), i.e. for each position of interest and each sample, a vector of frequency of amino acid alleles was determined by Anvi’o based on read mapping. For each TR-VR pair, all amino acid residues in the VR for which at least one of the position in the codon corresponded to an A in the TR were evaluated, along with 10 randomly chosen positions upstream in the target protein sequence which were used as control.

Three complementary metrics were computed from these amino acid alleles frequency vectors. First, the entropy calculated by Anvi’o was used as a measure of the populations diversity at a given position in a given sample. Based on the overall distribution of entropy values across VR and control positions, we established three categories of positions: low entropy for values ≤0.25, medium entropy for values >0.25 and ≤0.5, and high entropy for values >0.5. Next, we calculated for each position a cosine similarity between the allele frequency vector of a sample and the allele frequency vector of the previous sample in the time series. Again, based on the overall distribution of cosine similarities across consecutive time points for VR and control positions, we defined cosine similarity values ≥0.9 as “high similarity”, values ≥0.75 and <0.9 as “medium similarity”, and values <0.75 as “low similarity”. Finally, we calculated the number of changes in the dominant (i.e. majority) allele throughout the time series for each position.

Four categories of DGR activity were then defined based on a combination of these 3 metrics. First, if the entropy of position was always high, i.e. >0.5 for all time points, the position was considered as “constant diversity”, i.e. it is likely that the corresponding DGR is active enough to counteract any purifying selection. If instead a position included in the same time series included both samples with low (i.e. ≤0.25) and high (i.e. >0.5) entropy, it was considered as “alternating”, i.e. these changes were interpreted as a series of DGR-driven diversification events followed by diversity reduction through purifying selection and/or drift. Alternatively, if at least one dominant allele change was observed during the time series with a low cosine similarity (i.e. <0.75), the corresponding DGR was also considered as “alternating”. In this case, we interpreted the change in dominant allele associated with a high distance between allele frequency vectors as evidences suggesting some unsampled DGR diversification event between the time points. The cutoff on distance between allele frequency vectors enabled us to distinguish cases with genuine changes in the amino acid composition of a position, from cases with multiple co-dominant alleles of nearly equal proportion, for which a change in dominant allele could be observed by chance without the need for any diversification or selection event. Positions not considered as “constant diversity” or “alternating” were then classified as “constant selection” if the minimum entropy was at least medium (i.e. >0.25). These are cases for which few to no change in the dominant allele are observed, however all samples show significant population diversity suggesting a continuous DGR-driven diversity likely controlled by purifying selection. Finally, other positions with either all samples with entropy ≤0.25 (low entropy) or all similarities between samples ≥0.9 (high similarity) were considered as “inactive”. We chose to interpret these as sign that the corresponding DGR was not active, although similar allele frequency profiles could be obtained from active DGR associated with very strong purifying selection if the fitness of different variants was constant across the time series.

Overall, >97% of the “control” positions (i.e. positions from the target gene but not in the VR region) were classified as “inactive”, as would be expected for positions under strong purifying selection. In contrast, only 43% of the VR positions were classified as “inactive”, despite the fact that the VR regions were determined automatically and likely include non-VR positions in 5’ and 3’ of the actual TR-VR repeat. The other VR positions distributed between “constant diversity” (35%), “alternating” (15%), and “constant selection” (7%). This is consistent with the high rate of non-synonymous SNV identified from individual metagenome mapping (Fig. 3A & B), and confirms that the population diversity observed at VR locus is fundamentally different from the one observed at other positions even in the same target gene and the same samples.

### Estimation of the contribution of DGRs to overall amino acid changes in viral genomes

To conservatively estimate the contribution of DGRs to overall amino acid turnover in viral genomes, we first selected DGRs from clades 1, 4, and 6 for which longitudinal data was available, and excluded data from laboratory incubations (Supplementary Table 6). For each type of dataset (“Temperate_Lakes”, “Antarctic_Lakes”, “Human_microbiome”, “Human_microbiome_perturbed”), the total number of changes in dominant allele observed on VR and control positions (see above) was tallied and divided by the total number of observation for each group to obtain a “frequency of change” for each group. Based on an average genome size of ∼50kb and coding density of ∼90% for *Caudovirales* (based on genomes in NCBI Viral RefSeq v93), we estimated an average number of position per genome of 15,000 (45,000 nucleotide positions in protein-coding gene leading to 15,000 codons/amino acid residues). For VR positions, we used the average number of VR positions predicted by DGR, i.e. 17. For each dataset, the average frequency of change for each group (background and VR) was thus multiplied by the estimated number of positions for each group in an average genome (15,000 and 17) to obtain estimates of total number of changes for each group. The ratio between the number of changes in VR and the total number of changes was then used as an estimate of the contribution of DGRs to amino acid turnover in an average DGR-encoding viral genome.

The estimated proportions of amino acid changes associated with DGRs were 6.14% in “Human_microbiome”, 8.05% in “Human_microbiome_perturbed”, 9.67% in “Temperate_Lakes”, and 16.35% in “Antarctic_Lakes”. Importantly, we consider these estimates as conservative because all positions randomly selected as ‘control’ were taken from outside the predicted VR but within the same DGR target gene. These genes are likely to experience more frequent changes even outside of the VR region than other housekeeping genes because most of them are directly involved into virus-host interactions. Hence, these estimated proportion of DGR-driven amino acid changes should be seen as lower boundaries of the actual value.

