## Supplementary Figures for "Ecology and molecular targets of hypermutation in the global microbiome"

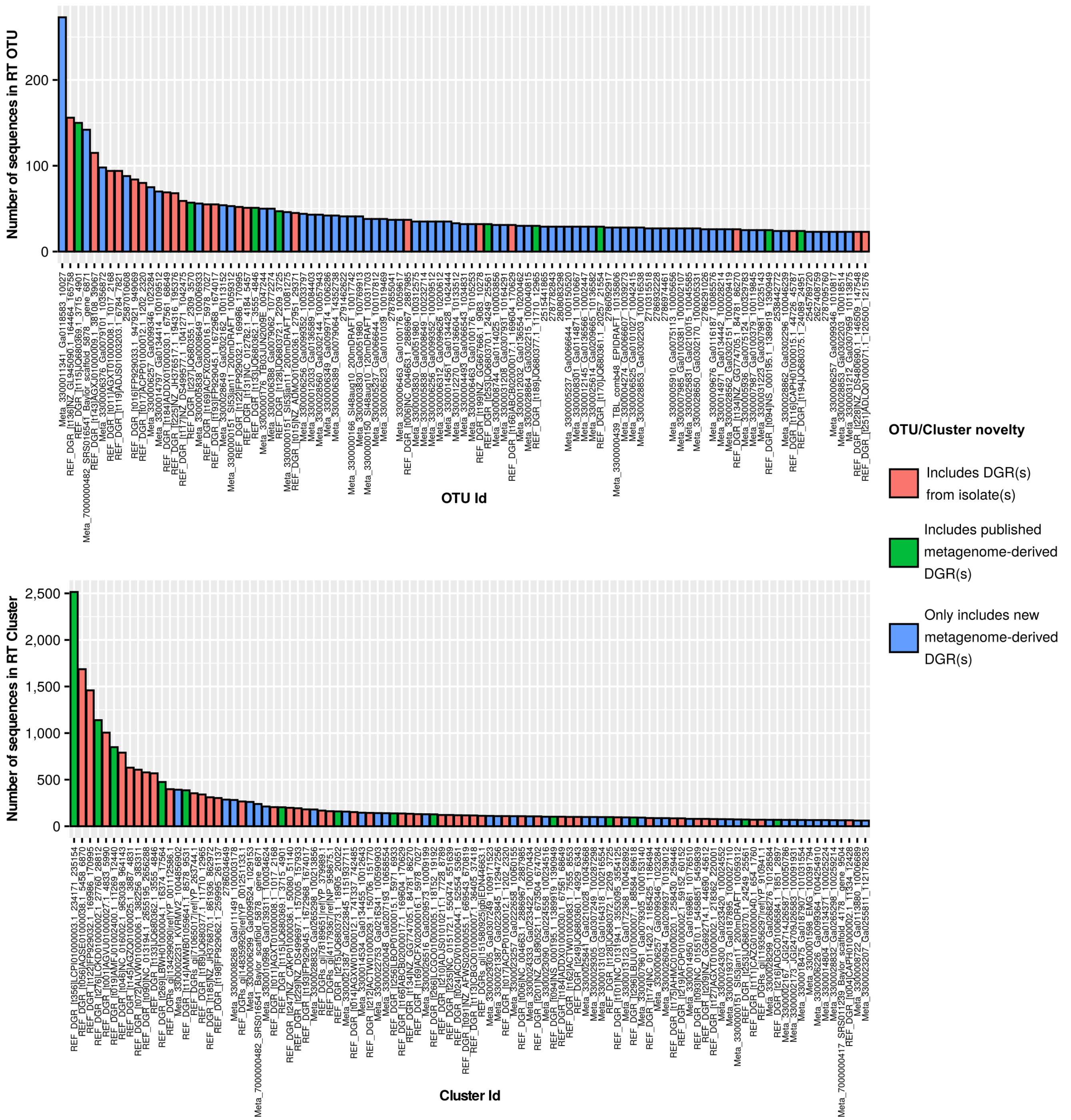


**Supplementary Figure S1: Size distribution of the 100 largest RT OTUs (top panel) and RT Clusters (bottom panel).** The bars are colored according to the presence/absence of reference sequences in the corresponding OTU/Cluster.

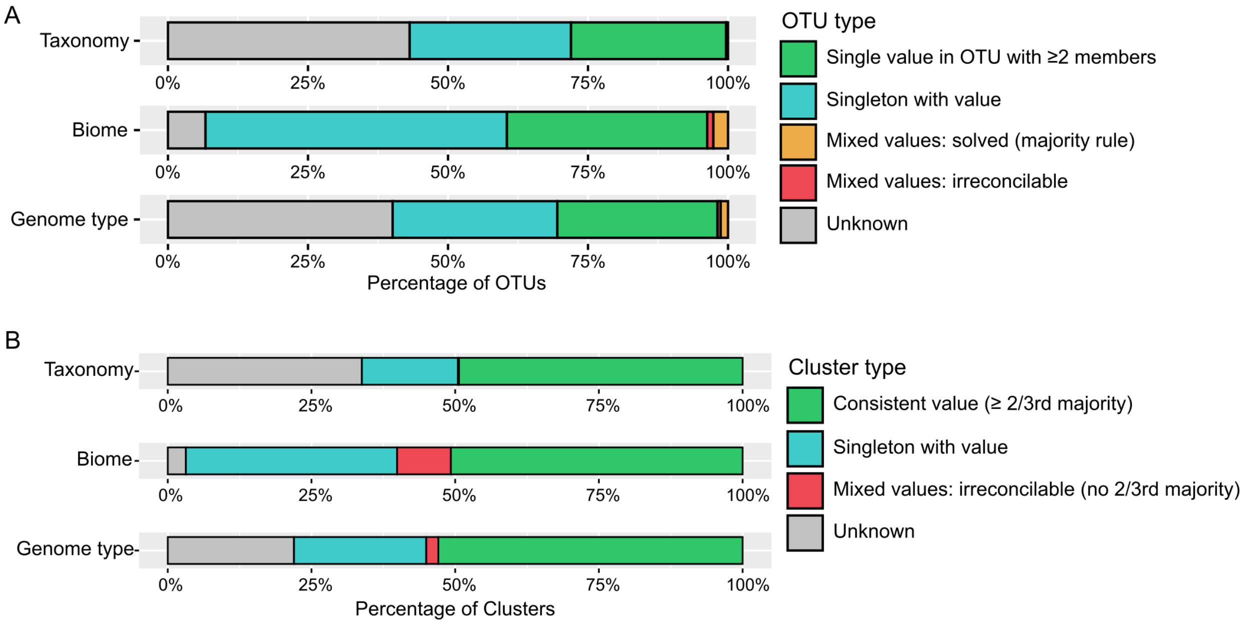


**Supplementary Figure S2: Characteristics of DGR RT OTUs and Clusters. Each bar chart indicates the consistency of one feature (taxonomic classification, biome, or genome type) across members of a DGR RT OTU (A) or DGR RT Cluster (B).** For OTUs with ≥2 members (i.e. non-singletons) with inconsistent values for a feature, a majority rule was applied: if a majority of an OTU members had the same value, this value was used for the OTU ("Mixed values: solved"). In case of tie (i.e. equal number of members associated to different features), the OTU feature was considered as unknown ("Mixed values: irreconcilable"). For Clusters, a similar approach was used with a 2/3rd majority rule. All Clusters for which ≥ 2/3rd of the members had the same value were considered as "Consistent value" and the value was assigned to the cluster. Cases in which the majority value in the cluster was associated with < 2/3rd of the members were considered as "Mixed values: irreconcilable".

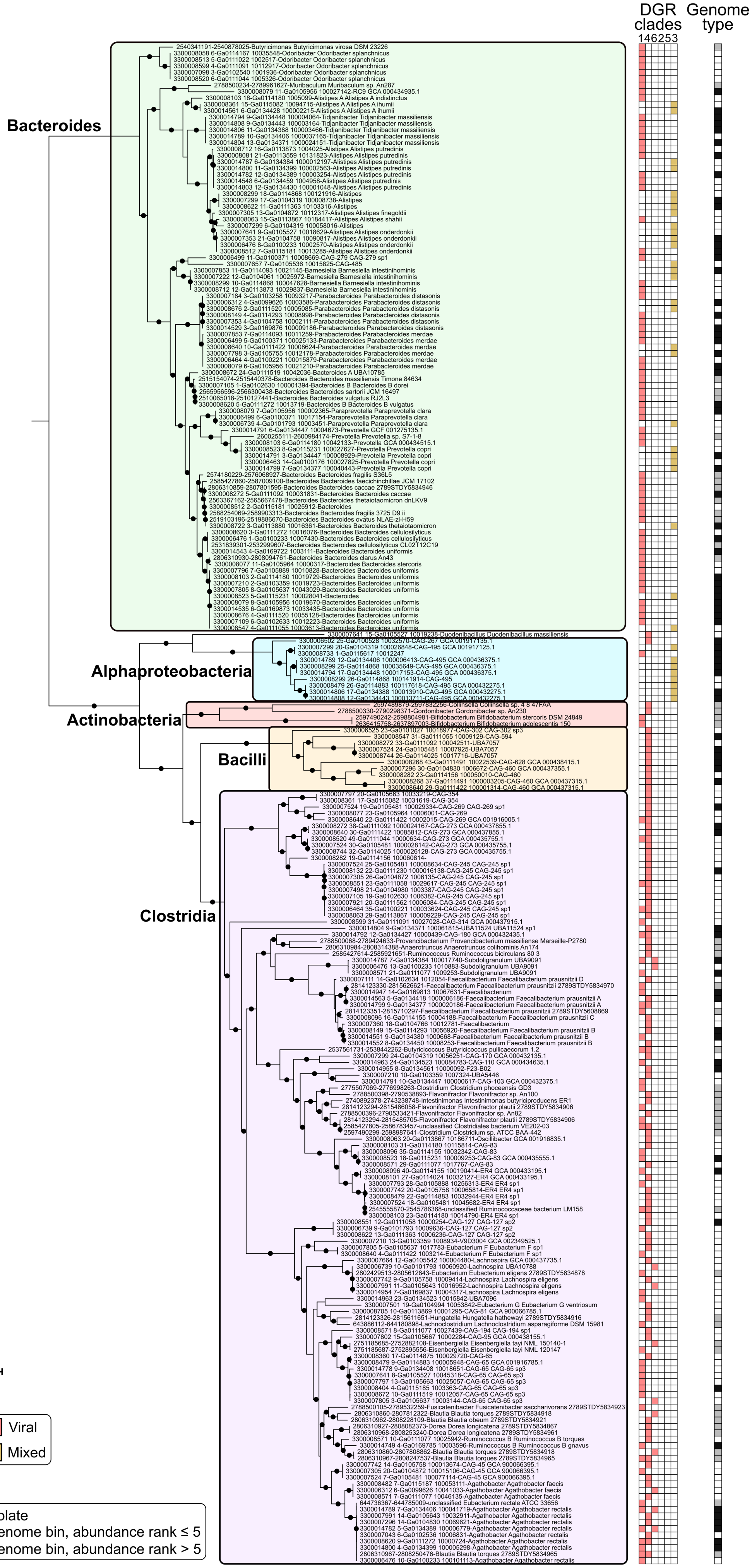

DGR clades
1 4 6 2 5 3
Genome type
Bacteroides
Alphaproteobacteria
Actinobacteria
Bacilli
Clostridia
Tree scale: 0.1
DGR clade type
Viral
Mixed
Genome type
Isolate
Genome bin, abundance rank ≤ 5
Genome bin, abundance rank > 5

**Supplementary Figure S3: Phylogeny of isolate and metagenome-binned genomes encoding one or more DGRs and associated with human gut samples.** Nodes with support <50 were collapsed, and nodes with support ≥80 are noted with a black circle. For each genome, the different clades of DGRs detected in the genome is indicated next to the tree as a colored heatmap. The genome relative abundance is then indicated next to the heatmap: isolate genomes are highlighted with grey squares, genome bins ranked as one of the 5 most abundant genomes within a metagenome are highlighted with black squares.

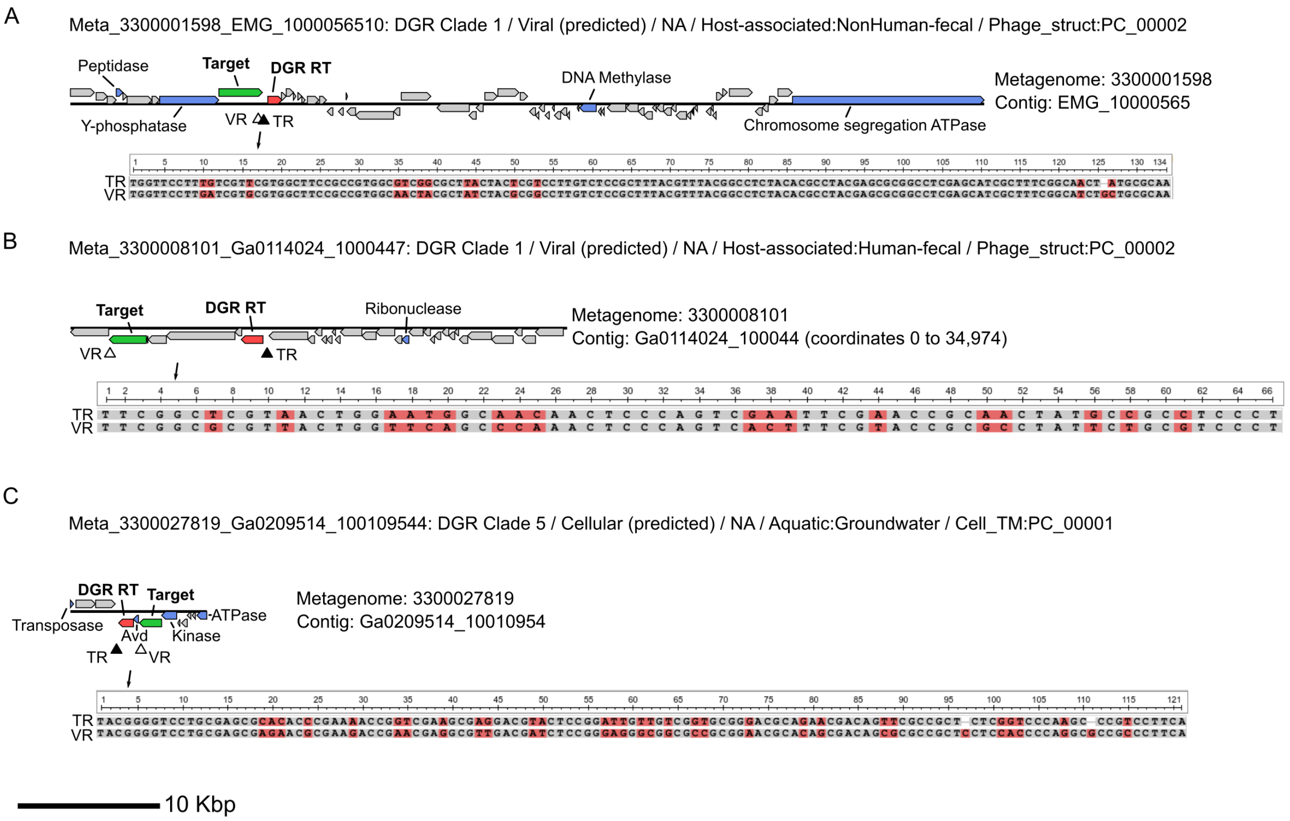


**Supplementary Figure S4: Examples of predicted DGRs with atypical (non-A) mutation bias.** For each DGR, the clade, genome type, taxonomic classification, biome, and primary target affiliation are indicated when available. The genome maps are colored based on each predicted CDS functional annotation: the DGR reverse-transcriptase in red, target gene in green, other genes in blue, and “hypothetical protein” in grey.

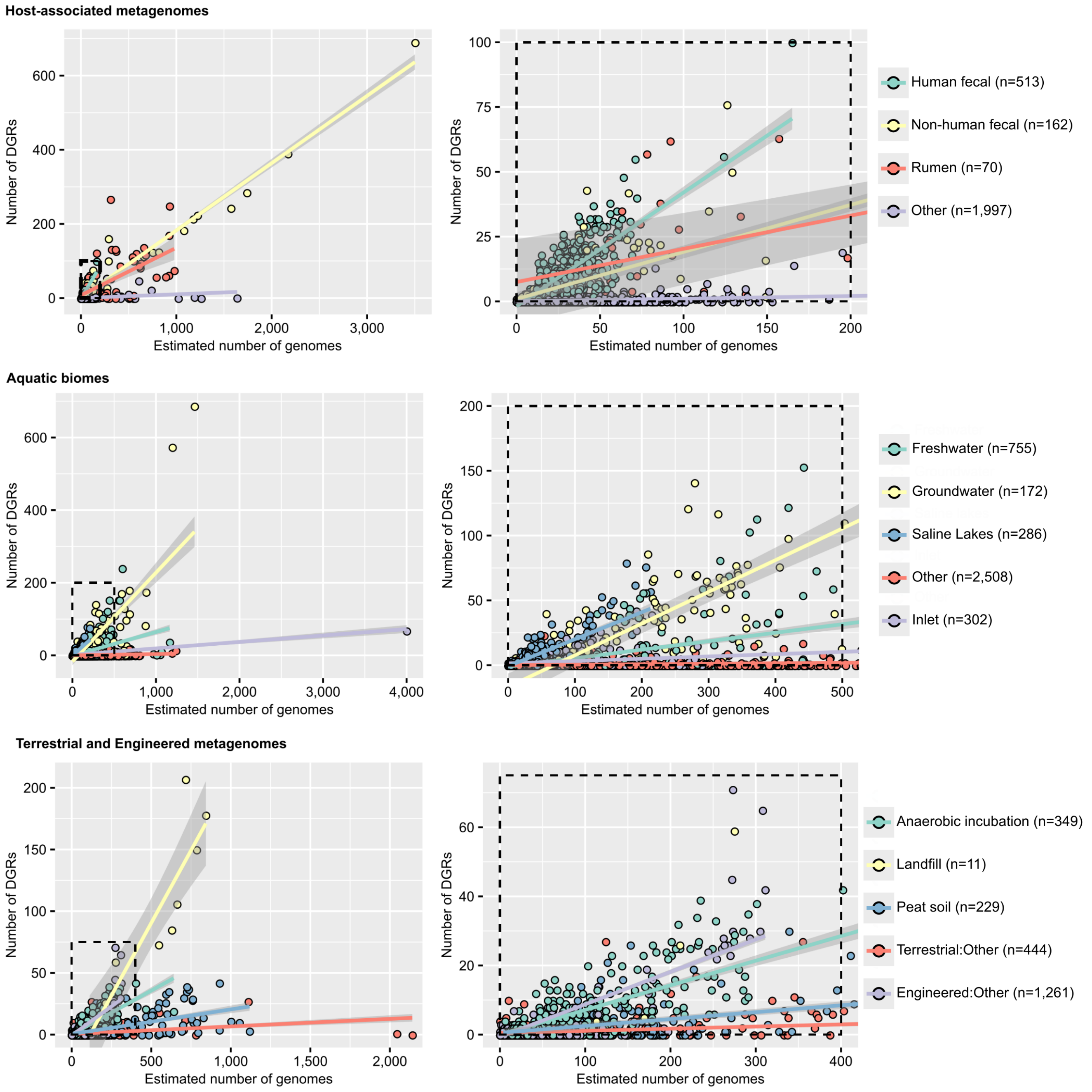


**Supplementary Figure S5: Link between estimated total number of genomes (x-axis) and number of DGRs detected (y-axis) for metagenomes across different biomes.** For each biome, a linear regression line is indicated in color, with the 95% confidence interval outlined in gray. Zoomed plots are displayed on the right panel, and the zoomed-in region is highlighted with a dashed black square on the full plot on the left panel.

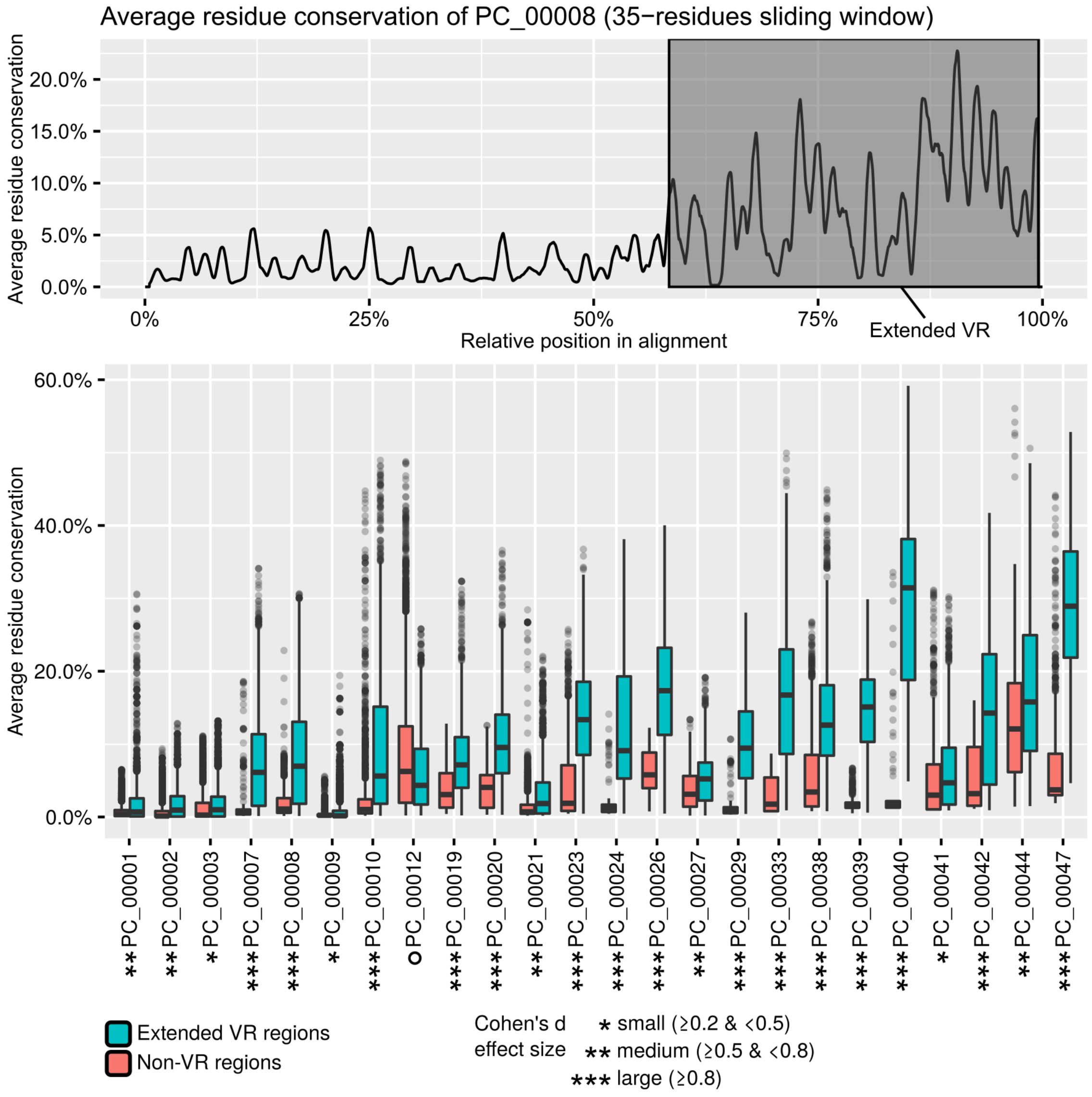


**Supplementary Figure S6: Average residue conservation in predicted targets. A. Example of average residue conservation in 35-residues windows along the multiple alignment of PC_00008.** An "extended" VR region (200 residues upstream and 20 residues downstream of the average predicted VR region) is highlighted in grey, which corresponds to the variable residues and the surrounding conserved domain. B. Distribution of residue conservation in "extended VR" and non-VR regions for the 24 largest target clusters. All distribution were significantly different (Kolmogorov–Smirnov test p-value <2E-16). The magnitude of the difference between VR and non-VR region is indicated through Cohen's d effect size (star symbols on the x-axis). All target PCs showed a higher average conservation in VR compared to non-VR regions except for PC_00012, which is highlighted with a black circle. The boxplot lower and upper hinges correspond to the first and third quartiles, respectively, and the whiskers extend no further than ±1.5 times the interquartile range.

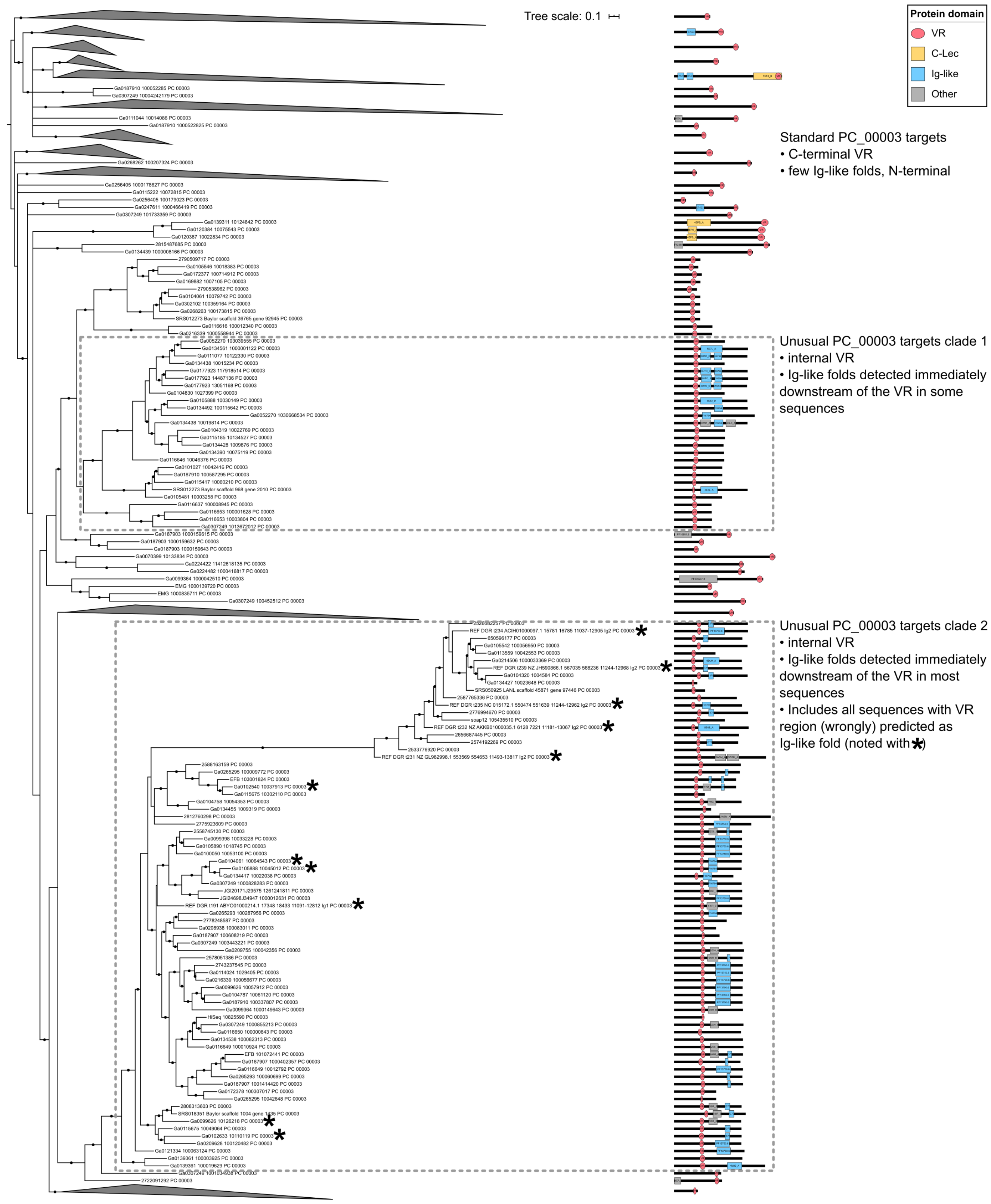


**Supplementary Figure S7: Phylogeny and domain organization of target sequences clustered in PC_00003.** Nodes with support <50 were collapsed, and nodes with support ≥80 are indicated with a black circle. For each sequence or clade, a schematic of the domain organization is indicated to the right of the tree, with a black line proportional to the sequence length, VR domains indicated with a red circle, and other domains indicated with colored rectangles. Monophyletic clades with a consistent domain organization were collapsed. The two clades of sequences displaying an internal VR region typically followed by one or several Ig-like folds in C-terminal are highlighted with a black dashed square. Reference sequences identified in the “Ig1” and “Ig2” domains are noted with a star symbol next to the sequence name.

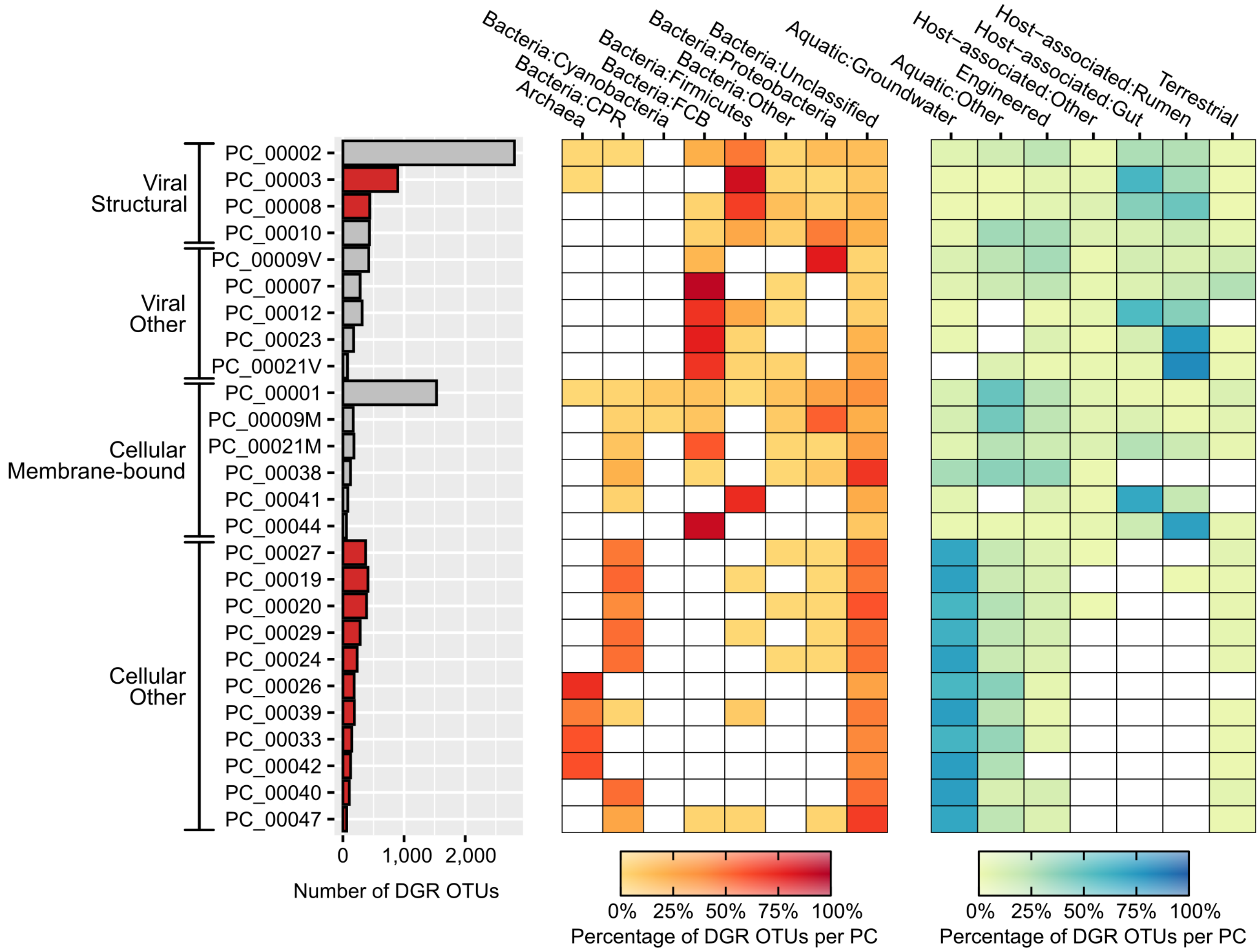


**Supplementary Figure S8: Taxonomic classification and biome of DGR OTUs associated with the 24 largest target PCs.** The PCs are ordered according to the 4 main categories of targets, as on Fig. 2. Target PCs for which the VR region was not identified as a putative C-Lec fold are highlighted in red. The proportion of DGR OTUs associated with specific taxa (left) or biomes (right) was calculated independently for each target PC. White cells in the heatmap correspond to an absence of DGR for the corresponding taxa/biome and target PC combination. CPR: Candidate Phyla Radiation. FCB: Flavobacteria, Fibrobacteres, Chlorobi, Bacteroides.

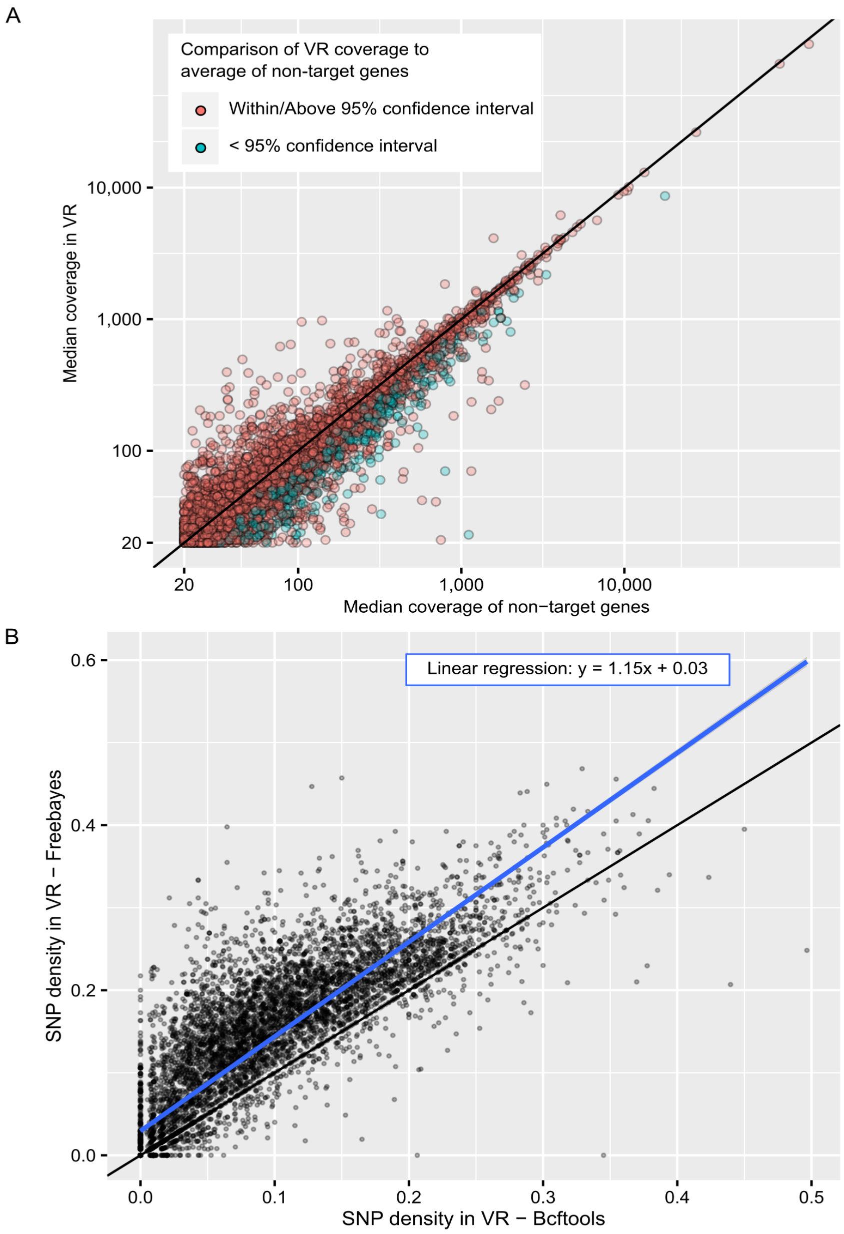


**Supplementary Figure S9: Read mapping and SNV calling on VR regions.** A. Comparison of coverage between VR regions and non-target genes for individual TR-VR pairs. Only cases with coverage ≥20x are displayed, and both x- and y-axis are displayed as log10 scale. The 1-to-1 line is indicated in black. A lower bound for a 95% confidence interval was calculated from the average coverage of non-target genes from the same contig minus 2 standard deviations. If the VR coverage was below this cutoff, it was considered as significantly lower than expected, the TR-VR was colored in blue in this plot, and flagged as "low coverage" if no SNVs were detected in Fig. 3A. B. Comparison of SNV density for individual VR regions obtained from Mpileup (x-axis) vs Freebayes (y-axis). A linear regression curve is plotted in blue, and the associated equation is indicated on the plot (p-value <2e-16).

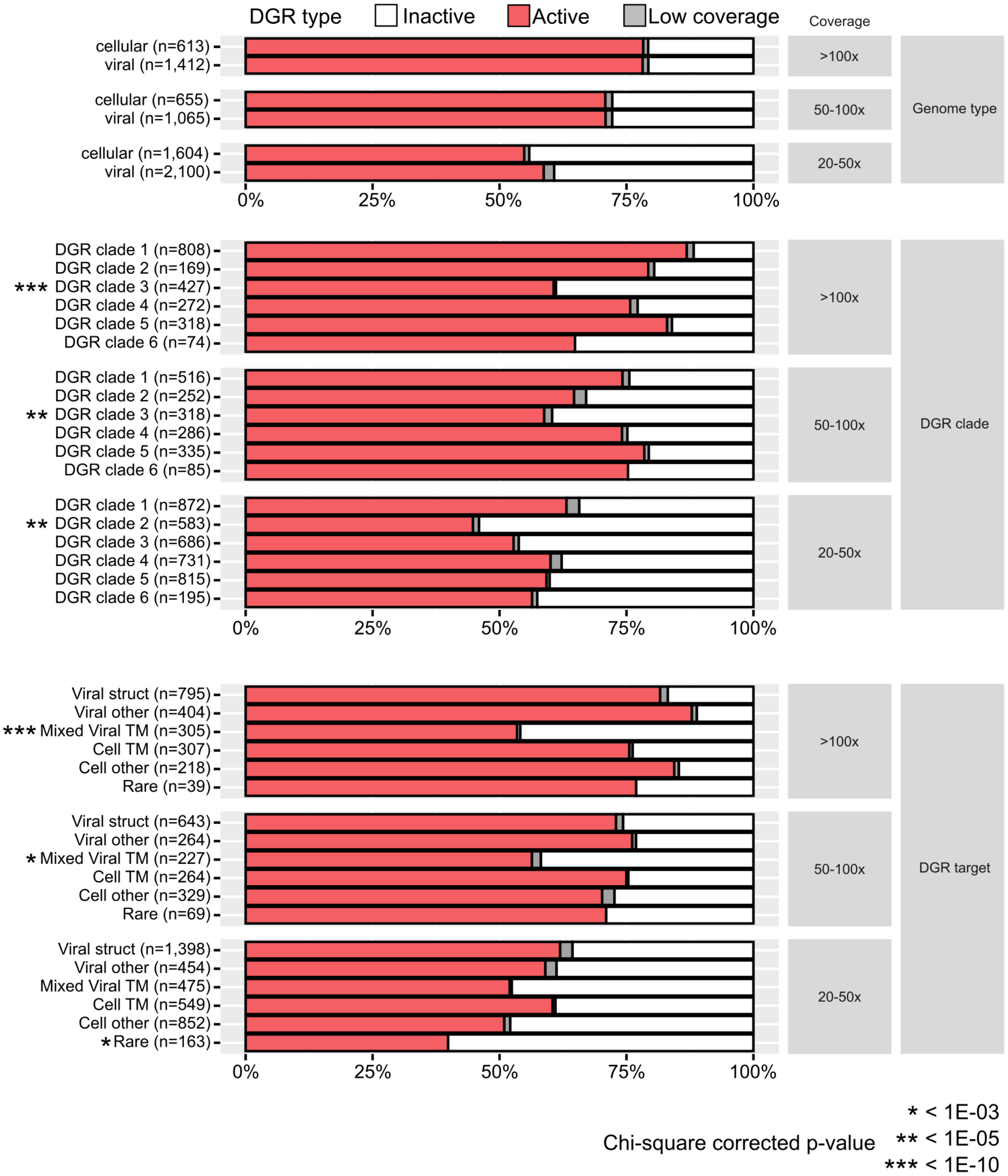


**Supplementary Figure S10: Distribution of active-vs-inactive DGRs across genome type, clade, and targets, for different ranges of coverage.** Groups (i.e. DGRs of the same genome type, DGR clade, or target) with a significantly lower proportion of active sequences compared to the average of the corresponding coverage category (Chi-squared test of independence) are highlighted with star symbols (Bonferroni-corrected p-values: *<1E-03, **<1E-05, ***<1E-10).

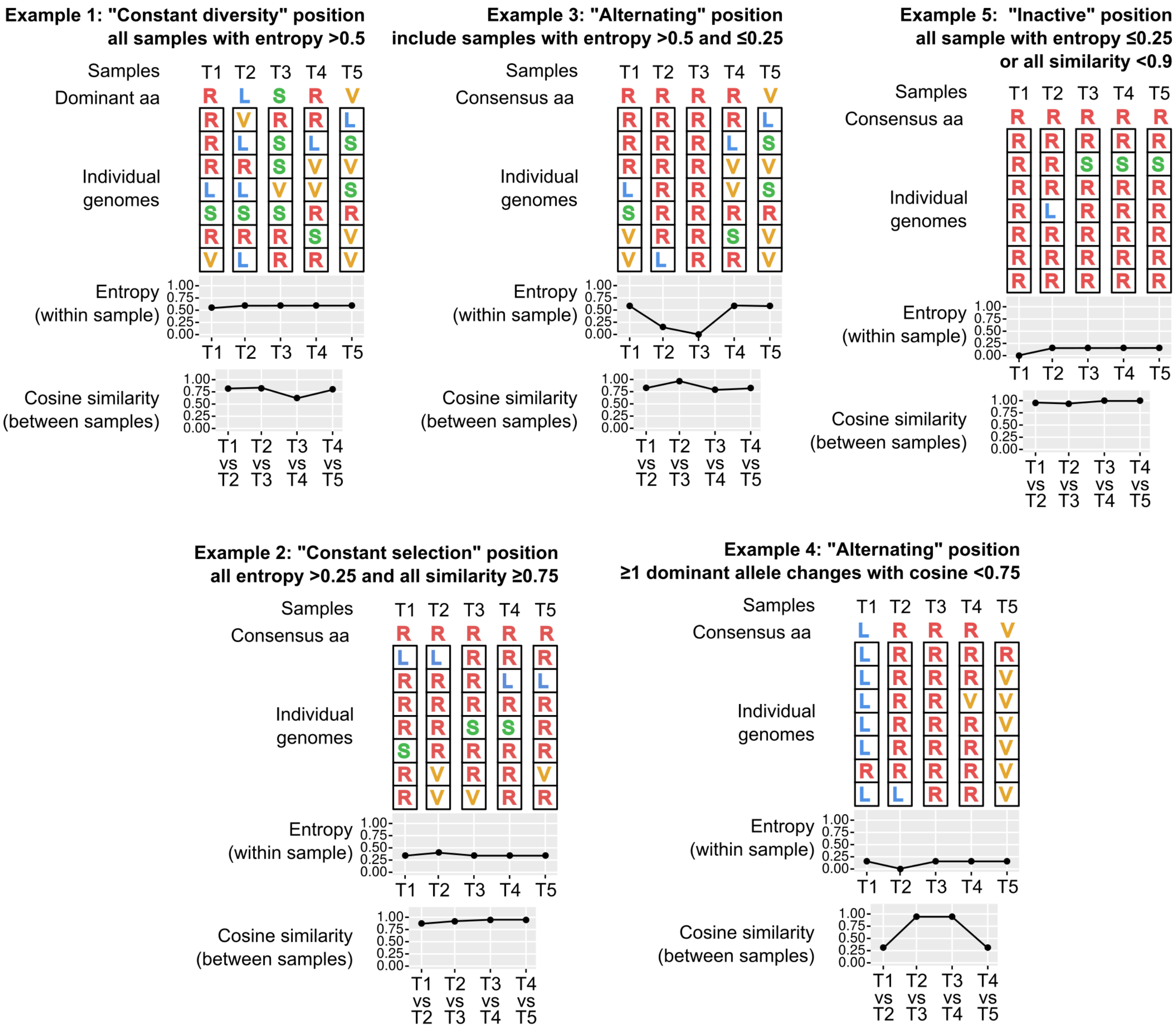


**Supplementary Figure S11: Schematic of the different categories of positions defined based on population diversity across time series.** Each example represents an individual position observed across 5 samples. The population diversity in each sample is represented as a heatmap, and the two metrics used to define the DGR activity categories are plotted underneath, either for each sample for the entropy, or between pairs of consecutive samples for the cosine similarity.

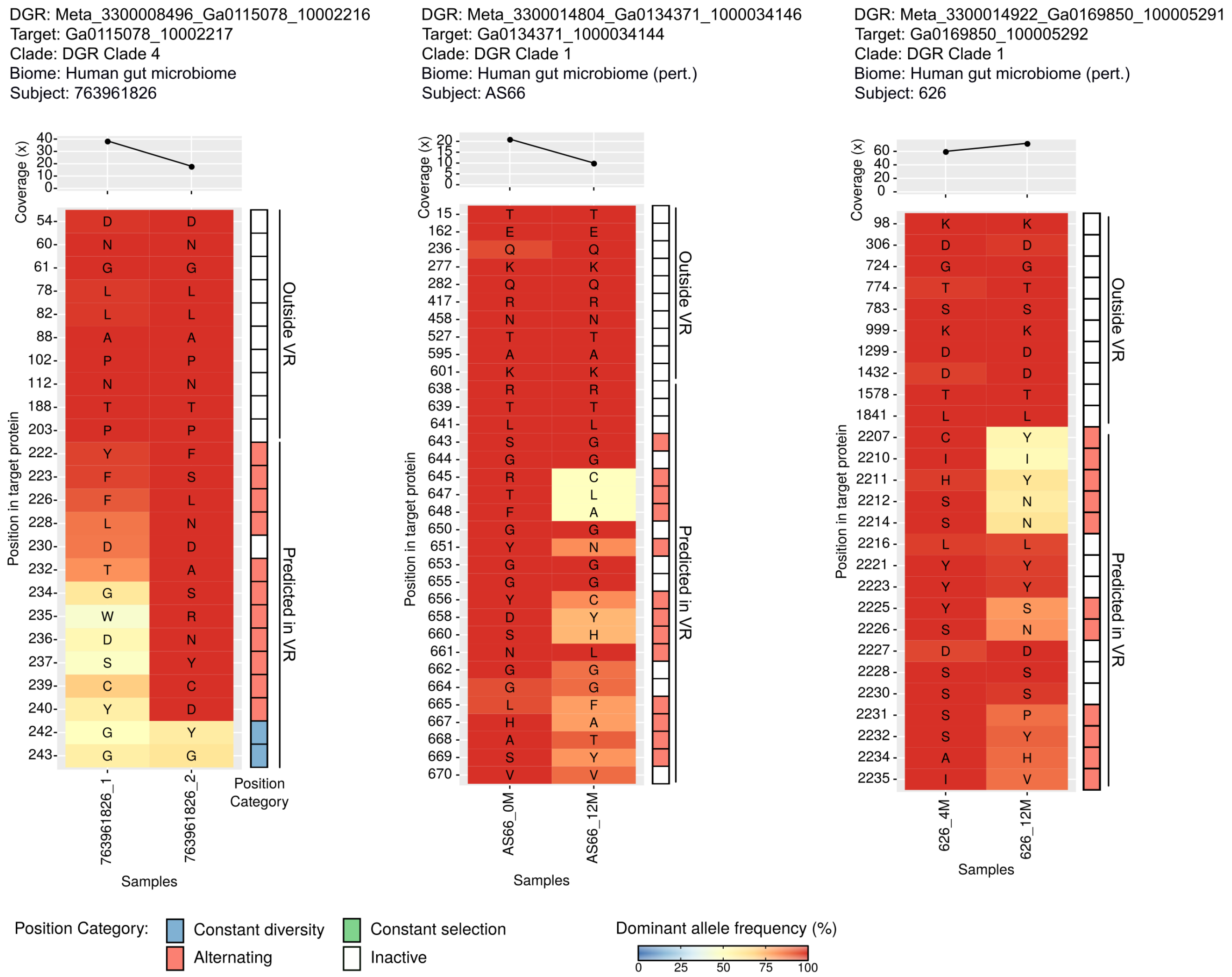


**Supplementary Figure S12: Examples of DGR target positions with changes in dominant amino acid between samples and low diversity within sample ("Alternating" pattern in Fig. 3D).** For each position (y-axis), the corresponding amino acid is indicated in the main heatmap with its frequency within the population indicated in color for each sample (x-axis). The right panel indicates the category of the position based on within-sample entropy, between-samples cosine distances, and number of amino acid changes in the time series (see Supplementary Text), colored as in Fig. 3D. The top panel indicates the median coverage of all positions in each sample. For reference purposes, 10 random positions from the same protein but outside of the predicted VR are included in the heatmap.

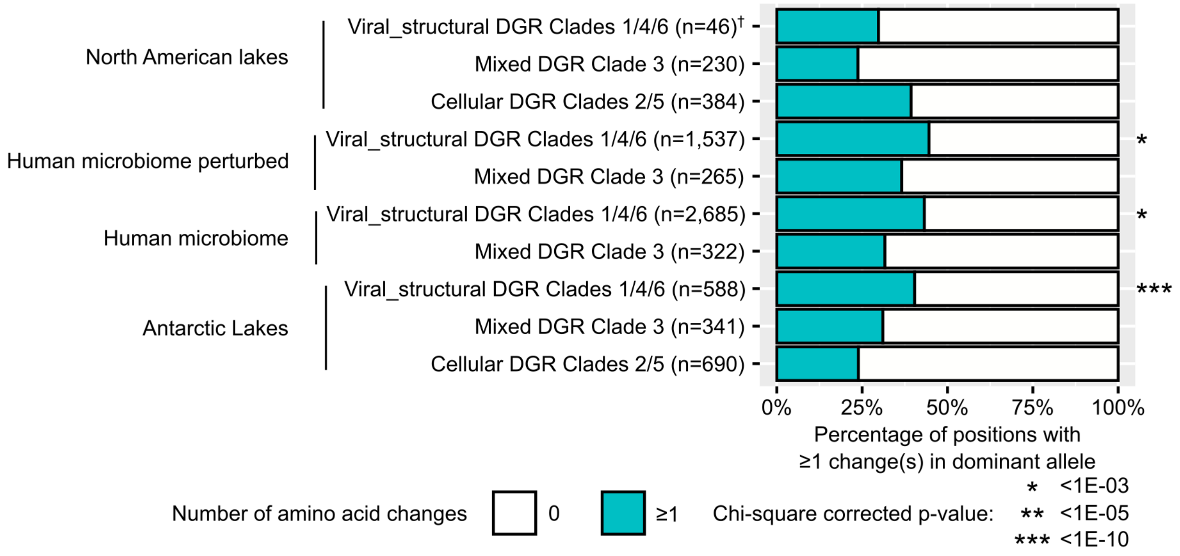


**Supplementary Figure S13: Percentage of positions with ≥1 change(s) in dominant allele among positions considered as "Constant diversity" or "Alternating" for different types of DGR across major biomes.** For each biome, the percentage in "Viral structural – Clades 1/4/6" DGRs was compared to the percentage in other DGR categories combined using a Chi-square test of independence. Groups with a significantly higher proportion of positions with ≥1 change(s) are highlighted with star symbols (Bonferroni-corrected p-values: *<1E-03, **<1E-05, ***<1E-10). † Counts for DGRs associated with viral structural proteins in temperate lakes are based on only 5 DGRs, while all other environments had > 10 DGRs associated with viral structural proteins.

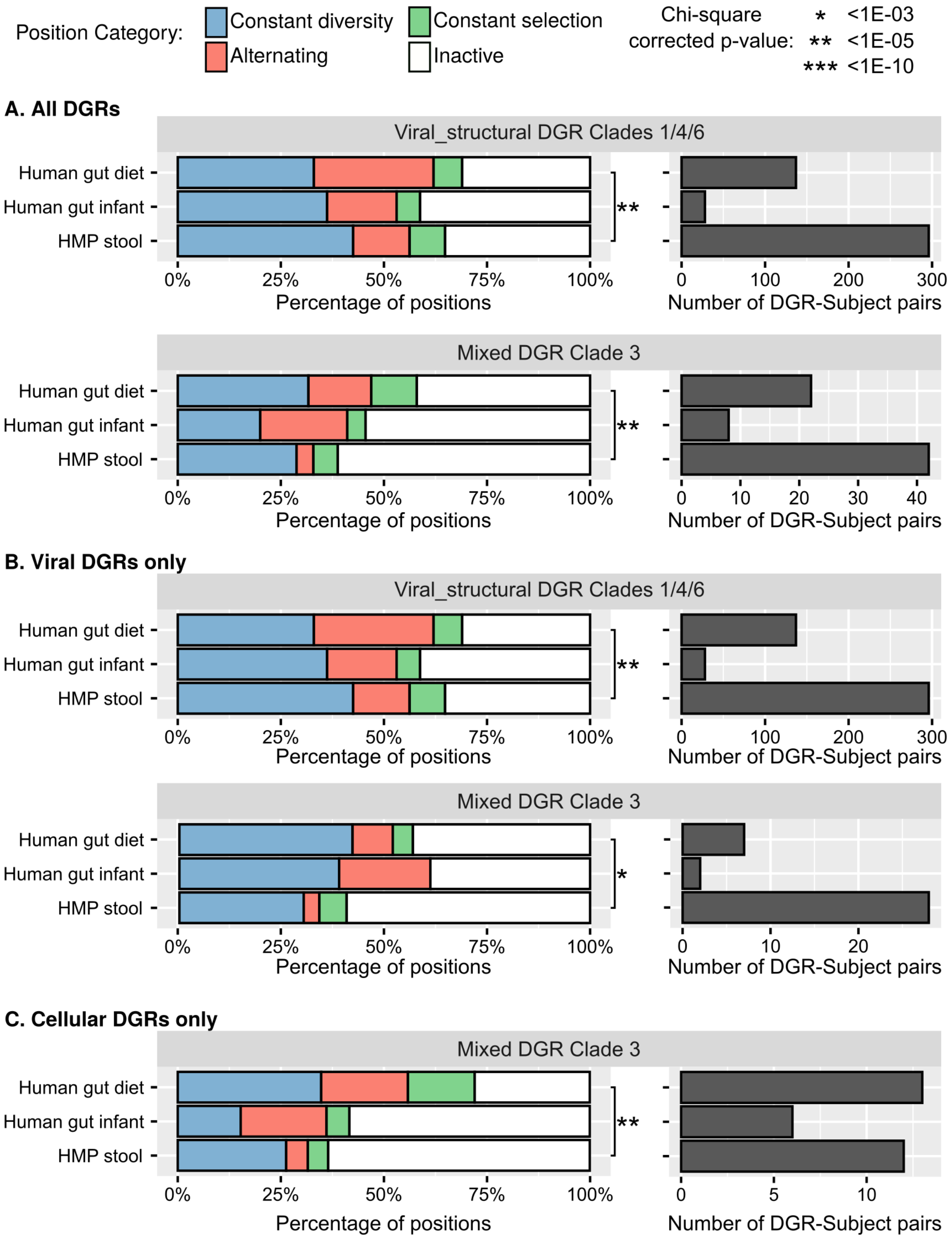


**Longitudinal datasets**
Human gut diet: 15 adults, 52-week weight-loss program, up to 5 time points (perturbed)
Human gut infant: 14 infants, 1st year of life, up to 3 time points (perturbed)
HMP stool: 40 adults, HMP production phase with multiple visits, up to 3 times points (control)

**Supplementary Figure S14: Comparison of DGR activity between perturbed and non-perturbed human microbiome samples.** Left panels display the distribution of activity categories for VR positions between perturbed and non-perturbed human gut microbiome DGRs. The conditions under which each dataset was collected are indicated at the bottom of the figure. The right panel bar graph indicates the number of observations (i.e. total number of DGRs covered in at least 2 time points across all subjects) for each dataset. Panel A includes all relevant DGRs, while panel B and C include only viral- or cellular-encoded DGRs, respectively. Statistically significant comparisons (Chi-square of independence) are highlighted with star symbols (Bonferroni-corrected p-values: *<1E-03, **<1E-05, ***<1E-10).
